# A mutant allele of the long-day flowering promoter NtFT5 unravels the mystery behind the short-day-specific flowering of tobacco cultivar Maryland Mammoth

**DOI:** 10.1101/2022.06.27.497884

**Authors:** Lena Grundmann, Marius M. Zimmermann, Andrea Känel, Axel Schwarze, David R. Wiedmann, Jost Muth, Richard M. Twyman, Dirk Prüfer, Gundula A. Noll

**Author notes:** These authors have contributed equally to this work (Lena Grundmann, Marius M. Zimmermann). For correspondence: Gundula A. Noll; Schlossplatz 8, 48143 Münster. M. Zimmermann, A. Schwarze and D. Wiedmann were affiliated with UM IBBP/Fraunhofer IME at the time the work was completed. Authors’ email addresses.

## Abstract

Flowering in day-neutral tobacco (*Nicotiana tabacum*) plants requires the photoperiod-dependent expression of members of the FLOWERING LOCUS T (FT)-like clade of phosphatidylethanolamine-binding proteins. FT-like floral activators and inhibitors compete for interaction with FD proteins to shift from vegetative to reproductive growth. In the short-day (SD) cultivar Maryland Mammoth (MM), vegetative growth persists under long-day (LD) conditions, generating unusually tall plants. We found that the major floral inducer under long-days (NtFT5) was expressed in MM and that *NtFT5* overexpression induced flowering in MM plants under LD conditions. However, sequence analysis revealed a 2-bp deletion near the 3⍰ end of *NtFT5* in MM plants resulting in a frame shift which leads to an altered amino acid sequence and a premature stop codon. We found that the truncated NtFT5_MM_ protein was still able to interact with tobacco FD proteins. However, constitutive overexpression under LD conditions in SD-specific flowering tobacco plants showed that NtFT5_MM_ is a weaker floral inducer than NtFT5. Our data suggest that the truncation does not impair the stability of the NtFT5_MM_ protein but may affect its binding affinity for NtFD1, probably resulting in the weaker expression of target genes. Our results therefore provide a potential explanation for the MM gigantism phenotype first observed more than 100 years ago.

**Highlight:** The previously unexplained gigantism of Maryland Mammoth tobacco is caused by a truncated major floral activator protein that results in weaker activation and the inability to flower under long-day conditions.

## Introduction

The transition from vegetative to reproductive development in flowering plants requires a precisely timed response to multiple endogenous and exogenous factors. In many plant species, a certain day length is required for floral induction. This perception of light and dark, and the subsequent adaptation, is known as photoperiodism, and plants can be assigned to one of three groups on this basis: short-day (SD), long-day (LD) and day-neutral. One of the first experiments addressing photoperiodism involved the tobacco (*Nicotiana tabacum*) cultivar Maryland Mammoth (MM), which does not flower under LD conditions and therefore grows unusually tall (Garner and Allard, 1920). Normally, tetraploid N. tabacum displays day-neutral flowering behavior. Many theories were proposed to explain the unanticipated obligate SD flowering of MM (including poor nutrition and transplant shock) before experiments confirmed that photoperiodic conditions were responsible for the LD-specific gigantism phenotype (Garner and Allard, 1920).

Since these early studies, the regulation of photoperiodic flowering has been investigated in a number of plant species, in many cases revealing that permissive photoperiodic conditions generate a mobile signal (florigen) that mediates the switch from vegetative to reproductive growth (for review, see Wickland and Hanzawa, 2015). It is now accepted that the FLOWERING LOCUS T (FT)-like protein family is a major part of that signal (for review, see Turck *et al*., 2008). In many plants, FT is produced in the leaves and transported via the phloem to the shoot apical meristem (SAM), where it interacts with the bZIP transcription factor FD to activate the floral transition (Abe *et al*., 2005; Wigge *et al*., 2005; Corbesier *et al*., 2007; Tamaki *et al*., 2007).

FT belongs to the phosphatidylethanolamine-binding protein (PEBP) family of developmental regulators, which are conserved in all eukaryotic organisms (Schoentgen and Jollès, 1995; Bradley *et al*., 1996; Banfield *et al*., 1998; Karlgren *et al*., 2011). Whereas FT proteins are generally floral activators, PEBPs from the TERMINAL FLOWER1 (TFL1)-like clade are floral inhibitors that maintain the inflorescence meristem (Bradley *et al*., 1996; Kobayashi *et al*., 1999). Interestingly, the *FT* gene family in some plants has undergone expansion and neofunctionalization, with some members evolving into floral inhibitors (Pin *et al*., 2010; Harig *et al*., 2012; Wickland and Hanzawa, 2015). For example, two antagonistic FT homologs have been identified in sugar beet (*Beta vulgaris*), where BvFT1 acts as a floral inhibitor by repressing the floral activator *BvFT2* at the transcriptional level (Pin *et al*., 2010). In day-neutral tomato plants (*Solanum lycopersicum*), flowering is regulated by the expression of four *FT*-like genes encoding the floral inhibitors SlSP5G, SlSP5G2 and SlSP5G3, and the floral activator SlSP3D. The *SlSP3D* gene is expressed similarly under LD and SD conditions, whereas SlSP5G is predominantly expressed under LD conditions, and *SlSP5G2* and *SlSP5G3* are predominantly expressed under SD conditions. Flower development in day-neutral tomato is therefore regulated by the differential photoperiodic expression of multiple *FT*-like genes (Molinero-Rosales *et al*., 2004; Lifschitz *et al*., 2006; Cao *et al*., 2015).

The tobacco genome also encodes multiple FT homologs, some of which (NtFT1–NtFT3) are floral inhibitors whereas others (NtFT4 and NtFT5) are floral activators (Harig *et al*., 2012; Beinecke *et al*., 2018; Wang *et al*., 2018). *NtFT1–NtFT4* are expressed predominantly under SD conditions whereas *NtFT5* is expressed under both SD and LD conditions, and unlike *FT* genes from other species none of the *NtFT* genes show a circadian expression profile (Harig *et al*., 2012; Beinecke *et al*., 2018). Silencing the floral activator gene *NtFT5* by RNA interference significantly delayed flowering under LD conditions, whereas knocking out the NtFT5 gene using CRISPR/Cas9 rendered the mutants completely unable to flower under LD conditions, indicating that NtFT5 is a major floral inducer during long days (Beinecke *et al*., 2018; Schmidt *et al*., 2020). Three functional FD homologs have also been identified in tobacco (NtFD1, NtFD3 and NtFD4) and they interact with tobacco FT proteins (Beinecke *et al*., 2018). Furthermore, the *NtFT4* and *NtFT2* genes (encoding an activator and inhibitor, respectively) are expressed at similar levels under SD conditions, and the proteins show dose-dependent effects on flowering, which indicates they compete at the protein level for FD binding rather than using the mutual transcriptional regulation strategy described in sugar beet and potato (Pin *et al*., 2010; Abelenda *et al*., 2016; Beinecke *et al*., 2018). Indeed, NtFD1 preferentially interacts with the floral activator NtFT4 rather than the inhibitor NtFT2 (Beinecke *et al*., 2018). Taken together, these results show that although tobacco is a day-neutral plant, flowering is in part regulated by the photoperiod-dependent expression of different *FT* genes (Harig *et al*., 2012; Beinecke *et al*., 2018).

Here, we isolated and characterized the *NtFT5* gene in the SD cultivar MM and carried out overexpression studies to investigate its impact on flowering in MM compared to the day-neutral cultivar Hicks, which is considered to be isogenic to MM except at the MM locus (Smith and McDaniel, 1992). Our results provide insight into the molecular basis of a phenomenon that has been observed but not understood for more than a century (Amasino, 2013).

## Materials and Methods

### Plant materials and cultivation conditions

Seeds of *N. tabacum* cv. Hicks, *N. tabacum* cv. Maryland Mammoth (IPK Gatersleben, Germany) and *N. tabacum* SR1Δ*NtFT5* were sown in soil and cultivated in the greenhouse under LD conditions (16-h photoperiod, artificial light switched on if natural light fell below 700 μmol m^-2^ s^-1^, 22–25 °C under light, 20–25 °C in the dark) or in phytotrons under SD conditions (8-h photoperiod, 200 μmol m^-2^ s^-1^, 25 °C under light, 22 °C in the dark). Medial leaves were harvested 6 and 10 weeks after seed sowing (WASS) under LD and SD conditions and each sample comprised tissue pooled from three plants. Transgenic tobacco plants were generated by leaf disc transformation (Horsch *et al*., 1985) using *Agrobacterium tumefaciens* strain LBA4404 (Hoekema *et al*., 1983) and were regenerated on MS medium (Murashige and Skoog, 1962) containing 25 mg/L hygromycin for sterile selection under LD conditions (100 μmol m^-2^ s^-1^, 23 °C). After root formation, independent transgenic lines were transferred to the greenhouse and grown in soil. T_1_ seedlings were cultivated under sterile conditions on selective medium as above. Seedlings were either harvested for expression and western blot analysis or up to 10 individuals were transferred to the greenhouse for phenotypic analysis.

### Gene expression studies

Leaf or seedling samples were snap frozen in liquid nitrogen and stored at –80°C. Total RNA was extracted using the innuPREP Plant RNA kit (Analytik Jena, Jena, Germany) and residual genomic DNA was removed using the Turbo DNA-free kit (Thermo Fisher Scientific, Waltham, MA, USA). First-strand cDNA was synthesized by reverse transcription using PrimeScript RT Master Mix (Takara Bio Group, Saint-Germain-en-Laye, France). We carried out qPCR analysis using the CFX 96 Real-Time system (Bio-Rad, Munich, Germany) on a C1000 Touch Thermal Cycler. We mixed 0.1 µl template cDNA and 500 nM of each primer (Supplementary Table S1) with KAPA SYBR Fast qPCR Master Mix (Merck, Darmstadt, Germany). Each reaction comprised an initial denaturation step for 3 min at 95 °C, followed by 40 cycles of denaturation at 95 °C for 3 s and annealing/extension at primer-specific temperatures for 30 s (Supplementary Table S1). Melt curve analysis was carried out to confirm the specificity of the amplicon (5 s, 58–95 °C, ΔT = 0.5 °C). The target transcripts and reference *NtEF-1α* (Schmidt and Delaney, 2010) were analyzed in triplicate, and the quantification cycle (C_q_) values of technical replicates were averaged. The NTC (non-template control) and NRT (non-reverse-transcribed control) were analyzed in duplicate. Data were analyzed using Bio-Rad CFX Manager v3.1. The expression values of *NtFT4* and *NtFT5* were determined as previously described (Livak and Schmittgen, 2001).

### Cloning experiments

The *NtFT5* and *AtFT* coding sequences were amplified from tobacco cDNA (*N. tabacum* cv. Hicks and cv. MM) and *Arabidopsis thaliana* cDNA (cv. Col-0), respectively, using the primers listed in Supplementary Table S1. The overexpression constructs were prepared by digesting the NtFT5 and NtFT5_MM_ amplicons with the appropriate restriction enzymes and inserting them into vector pRT104 containing the CaMV 35S promoter and terminator (Töpfer *et al*., 1987). The entire expression cassette was then excised with HindIII and ligated into vector pBinHyg (Bevan, 1984). The expression constructs of *NtFT5* and *NtFT5*_MM_ under control of the endogenous P-NtFT5_2.6kb_ were prepared by digesting the P-NtFT5_2.6kb_, *NtFT5* and *NtFT5*_MM_ amplicons with the appropriate restriction enzymes and inserting them into vector plab12.1 containing the 35S terminator (linearized with XmaI/XbaI).

For the bimolecular fluorescence complementation (BiFC) assay, the *AtFT, NtFT5* and *NtFT5*_MM_ coding sequences were amplified using the primers listed in Supplementary Table S1, digested with the appropriate restriction enzymes and inserted into vector pENTR4 (Invitrogen, Karlsruhe, Germany). The pENTR4 constructs were then transferred to pBatTL NmRFP-*ccd*B by LR recombination using the Gateway LR Clonase Enzyme mix (Invitrogen). Split mRFP destination vectors containing *NtFD1, NtFD3, NtFD4* and NtFD1_SAAPF_ were already available (Beinecke *et al*., 2018). Destination vectors were kindly provided by Dr Guido Jach and Dr Joachim Uhrig, MPI Cologne, Germany (Müller *et al*., 2010). For the multicolor BiFC (mcBiFC) assay, vectors P35S:SCN-NtFT2:Tnos-Spacer and P35S:VN-NtFT2:Tnos pBluescriptII (Beinecke *et al*., 2018) were digested with StuI/BsrGI and ligated with the NtFT5 and NtFT5_MM_ coding sequences to generate vectors pBS-*SCN*-*NtFT5*-Sp, pBS-*SCN*-*NtFT5*_MM_-Sp, pBS-*VN*-*NtFT5* and pBS-*VN*-*NtFT5*_MM_. To generate dual expression cassettes, P35S:*VN-NtFT5*_(MM)_ was excised with ClaI/DraIII and transferred to pBS-*SCN*-*NtFT5*_(MM)_-Sp to create four intermediate vectors. The dual expression cassettes were excised using PmeI/BamHI and transferred to plab12.1 (linearized with XmnI/BamHI) to create four binary vectors named plab-*SCN*-*NtFTx*-Sp-*VN-NtFTx*. To generate the SCC-NtFD1 fusion, SCC-HA was amplified using the primers listed in Supplementary Table S1, digested with SalI/BamHI and transferred to pRT104 (linearized with XhoI/BamHI). The NtFD1 coding sequence was amplified, digested with BamHI/XbaI and transferred to pRT-SCC-HA (linearized with BamHI/XbaI). The P35S:SCC-NtFD1:T35S cassette was excised with HindIII and transferred to plab12.1 to create the binary vector plab-SCC-NtFD1.

The overexpression constructs for *NtFT5*_G170W_ and *NtFT5*_MM W170G_ were prepared by digesting the *NtFT5*_G170W_ and *NtFT5*_MM W170G_ amplicons (generated using the primers listed in Supplementary Table S1) with the appropriate restriction enzymes and inserting them into vector pRT104, containing the CaMV 35S promoter and terminator (Töpfer *et al*., 1987). The entire expression cassette was then excised with HindIII and transferred to vector plab12.10 (Post *et al*., 2014).

All constructs for transient and stable expression were introduced into the appropriate strain of *A. tumefaciens* by electroporation.

### Fragment length analysis

F_1_ and F_2_ individuals of the reciprocal cross of Hicks and MM were genotyped by fragment length analysis of the NtFT5 genomic sequence. This was carried out with a fragment of exon 4, containing the NtFT5_MM_-specific 2 bp deletion. The fragment was amplified using primers (see Supplementary Table S1) carrying the fluorescent dye 6-carboxyfluorescein (6-FAM). To identify NtFT5 and NtFT5_MM_ alleles the amplicon length of the NtFT5 allele from the cultivar Hicks and of the NtFT5_MM_ allele from the cultivar MM was simultaneously determined and used as references. The PCR samples were mixed with the GeneScan 600 LIZ dye Size Standard v2.0 (Thermo Fisher Scientific) and analyzed in an ABI 3730 Genetic Analyzer (Applied Biosystems, Waltham, MA, USA). The data were evaluated using the Applied Biosystems® Sizing Analysis Module Peak Scanner Software (version 3.0).

### Western blotting

Total protein extracted from 3-week-old transgenic and wild-type seedlings was boiled with Laemmli buffer (50% glycerol, 10% SDS, 0.25 M Tris-HCl pH 6.8, 5% 2-mercaptoethanol) containing 0.05% bromophenol blue and was separated by SDS-PAGE on a 12% polyacrylamide gel before electroblotting onto a nylon membrane. NtFT5 protein was detected using an anti-FT antibody (AS06198, Agrisera) diluted 1:1000.

### Grafting experiments

SR1 wild-type and 35S:*NtFT5*_(MM)_//SR1Δ*NtFT5* transgenic plants were cultivated in the greenhouse until floral buds became visible. Non-flowering SR1Δ*NtFT5* scions were then grafted onto each stock (n = 3) before further cultivation in the greenhouse. Emerging leaves of the SR1Δ*NtFT5* scions were removed regularly to enhance the source-sink gradient. To ensure seed production of P-NtFT5_2.6kb_:*NtFT5*//SR1Δ*NtFT5* and P-NtFT5_2.6kb_:*NtFT5*_MM_// SR1Δ*NtFT5* scions of these primary transformants were grafted onto florally induced SR1 wild-type plants.

### (Multicolor) bimolecular fluorescence complementation analysis

Transient expression was carried out by co-infiltrating the leaves of 3–4-week-old *N. benthamiana* plants (grown under LD conditions) with *A. tumefaciens* strain GV3101 pMP90 carrying the appropriate expression constructs for FT (*AtFT, NtFT5, NtFT5*_MM_, *NtFT5*_FN*_, *NtFT5*_FNC*_, *NtFT5*_R61G_ or *NtFT5*_R129A_) and FD (*NtFD1, NtFD1*_SAAPF_, *NtFD3* or *NtFD4*) and *A. tumefaciens* strain C58C1 carrying the pCH32 helper plasmid encoding virulence genes and the pBIN61 plasmid encoding the tomato bushy stunt virus RNA silencing suppressor protein p19 (Hamilton *et al*., 1996; Walter *et al*., 2004). The expression and helper strains were infiltrated at OD_600nm_ 1 and 0.3, respectively. We monitored mRFP fluorescence by confocal laser scanning microscopy (CLSM) using a TCS SP5 X microscope (Leica Microsystems, Wetzlar, Germany) at excitation and emission wavelengths of 549 and 569–629 nm, respectively. For mcBiFC (Hu and Kerppola, 2003; Waadt et al., 2008), the SCN/VN expression strains, the SCC-NtFD1 expression strain and the helper strain were infiltrated at OD_600 nm_ = 0.5, 0.33 and 0.3, respectively. Plants were cultivated under continuous light for 3–4 days, and leaf discs were screened for fluorescent cells in the abaxial epidermis. Fluorescence was observed by CLSM using a Leica STELLARIS 8 microscope (Leica Microsystems) at excitation/emission wavelengths of 458/470–490 nm for S(CFP) 3A and 488/505–525 nm for the VN/SCC chimera. Nuclear fluorescence was quantified using Leica LAS AF software and the data were analyzed using Origin 2020.

## Results

### Expression of endogenous floral activator genes *NtFT4* and *NtFT5* in SD cultivar MM and day-neutral cultivar Hicks

Flower development is induced under both LD and SD conditions in most tobacco cultivars, but cultivar MM flowers only under SD conditions (Fig. 1A–H). To determine whether the LD-specific non-flowering phenotype of MM reflects the silencing of *NtFT5*, the major floral activator under LD conditions (Harig *et al*., 2012; Beinecke *et al*., 2018; Schmidt et al., 2020), we compared the expression of *NtFT5* in MM and a day-neutral tobacco cultivar (Hicks) by quantitative real-time PCR (qPCR) (Fig. 1I–J). We found that NtFT5 was expressed in both cultivars under LD and SD conditions and the expression profiles were similar in each cultivar when matched for day length and the time point of harvest (6 or 10 WASS). *NtFT5* mRNA was generally less abundant under LD than SD conditions in both cultivars, and under SD conditions *NtFT5* was expressed at basal levels 6 WASS but was induced during floral transition (10 WASS). The main difference between the cultivars was that *NtFT5* mRNA levels were higher in Hicks than MM under LD conditions at the 6 WASS time point. However, *NtFT5* mRNA levels increased between 6 and 10 WASS under LD conditions in MM whereas we detected almost no difference in *NtFT5* mRNA levels between 6 and 10 WASS under LD conditions in Hicks. The increase in *NtFT5* mRNA levels between 6 and 10 WASS was less steep in MM under LD conditions than SD conditions. *NtFT5* mRNA levels were nearly identical in both cultivars 10 WASS under LD conditions, but Hicks flowered at this point whereas MM remained in the vegetative phase. In addition to *NtFT5* the closely related gene *NtFT7* was identified in whole-genome shotgun contigs of the *N. tabacum* cultivars K326, TN90 and Basma Xanthi, and it promoted the flowering of *N. sylvestris* plants constitutively overexpressing this gene (Beinecke *et al*., 2018). However, as previously described for cultivar SR1 (Beinecke *et al*., 2018), *NtFT7* was not detected in the genomes of cultivars Hicks or MM (Supplementary Fig. S1). We also analyzed the expression of *NtFT4*, which encodes a floral activator that is predominantly expressed under SD conditions in cultivar SR1 (Harig *et al*., 2012; Beinecke *et al*., 2018). Our qPCR results showed that *NtFT4* was also predominantly expressed under SD conditions in cultivars Hicks and MM, and the mRNA levels increased during the reproductive phase. *NtFT4* mRNA was also detected under LD conditions, mainly at 10 WASS, and the levels were comparable to those detected under SD conditions 6 WASS (Fig. 1I–J). Interestingly, flowering was delayed by a week in MM compared to Hicks under SD conditions, even though *NtFT4* and *NtFT5* were expressed at similar levels in both cultivars.

**Fig. 1.**
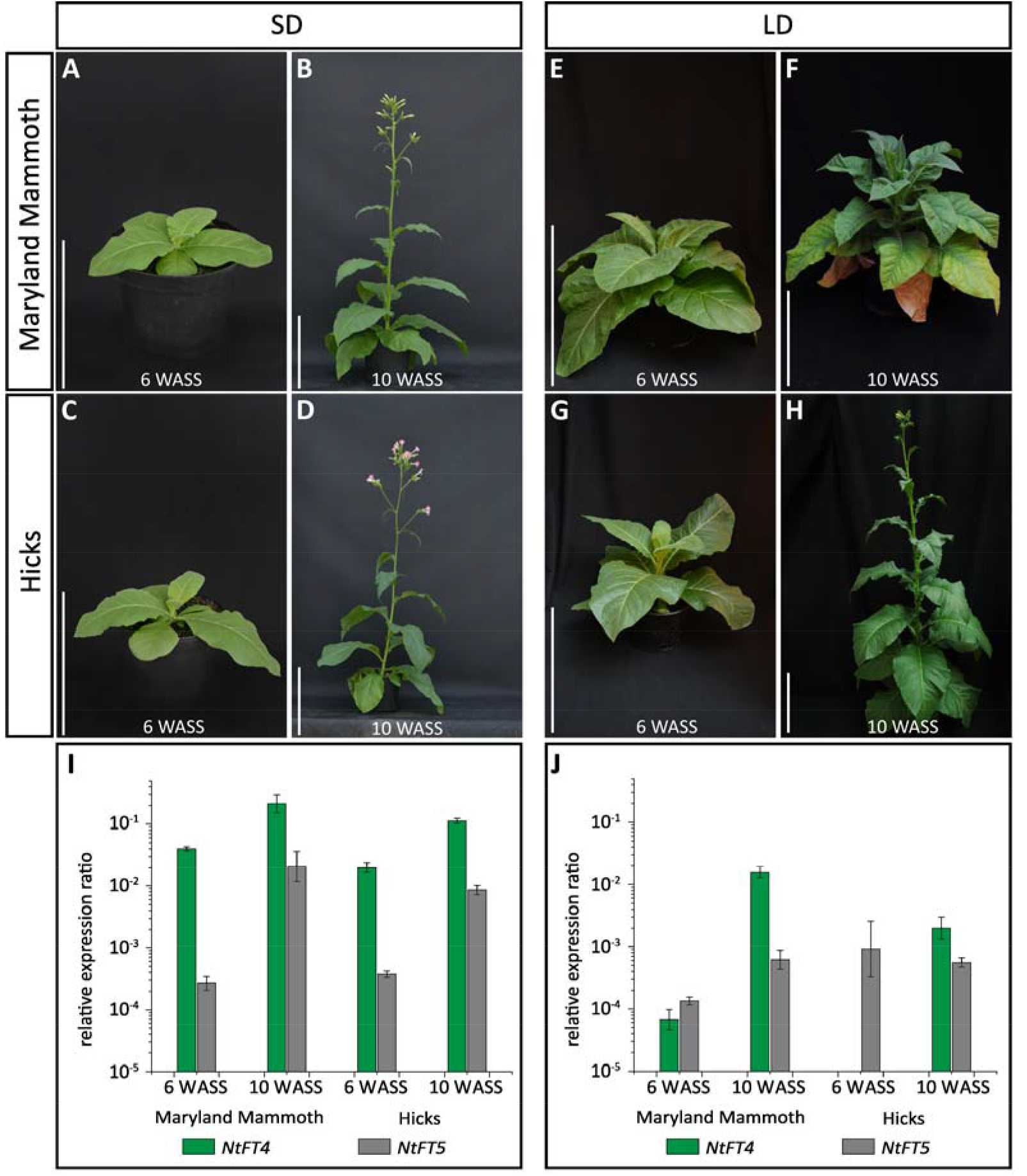
Phenotype and *NtFT4*/*NtFT5* gene expression in tobacco cultivars Maryland Mammoth (MM) and Hicks under short-day (SD) and long-day (LD) conditions. (A–D) Under SD conditions, flower development was observed both for (A,B) cv. MM and (C,D) cv. Hicks. (E–H) Under LD conditions, (E,F) no flower development was observed in cv. MM, but (G,H) flowers were produced by cv. Hicks. Pictures were taken 6 and 10 weeks after seed sowing (WASS). Scale bars = 30 cm. (I,J) *NtFT4* and *NtFT5* expression was analyzed by qPCR in three biological replicates per time point. Each biological replicate comprised pooled medial leaf tissue of three individual plants. Values were normalized to the reference gene *NtEF-1α*. Error bars represent the standard error of the mean of three biological replicates. *NtFT4* is predominantly expressed under SD conditions with rising mRNA levels during the reproductive phase in both cultivars. Under LD conditions, *NtFT4* mRNA levels were low (MM) or not detectable (Hicks) 6 WASS but increased in both cultivars 10 WASS. *NtFT5* expression increased with the initiation of floral development in both cultivars under SD conditions. Under LD conditions, the expression of *NtFT5* was higher in Hicks than in MM 6 WASS, but mRNA levels were similar in both cultivars 10 WASS.

### Floral induction in transgenic MM and Hicks plants expressing *NtFT5*

Having established that *NtFT5* is similarly expressed in both cultivars, we set out to determine whether there is a functional deficiency in the *NtFT5* gene in MM and whether MM has the propensity to flower under LD conditions when provided with a fully functional sequence. We therefore isolated the *NtFT5* coding sequence from Hicks (because this variety flowers under LD conditions) and transformed MM plants (and Hicks plants as a control) with the 35S:*NtFT5*_Hicks_ cassette. The constitutive expression of the *NtFT5*_Hicks_ gene caused a profound early flowering phenotype in both cultivars (Fig. 2A,B). We found that 50% of the lines began flowering in tissue culture and failed to form roots (four 35S:*NtFT5*_Hicks_ MM lines and five 35S:*NtFT5*_Hicks_ Hicks lines), but the other 50% could be transferred to the greenhouse and generated seeds. Expression analysis by qPCR revealed that transgenic lines from both cultivars accumulated ∼10^4^-fold more NtFT5 mRNA than the vector control (Fig. 2C,D). For a more detailed analysis, we studied the T_1_ progeny of two independent transgenic lines and one vector control line for each cultivar in the greenhouse under LD conditions (Fig. 2E,F). We observed a profound early flowering phenotype for all transgenic lines overexpressing *NtFT5*_Hicks_ resulting in significantly fewer leaves and significantly fewer days until flowering compared to the Hicks and non-flowering MM vector control lines. These results demonstrate that the *NtFT5*_Hicks_ gene can induce flowering during the early stages of development even under LD conditions when overexpressed in the SD cultivar MM.

**Fig. 2.**
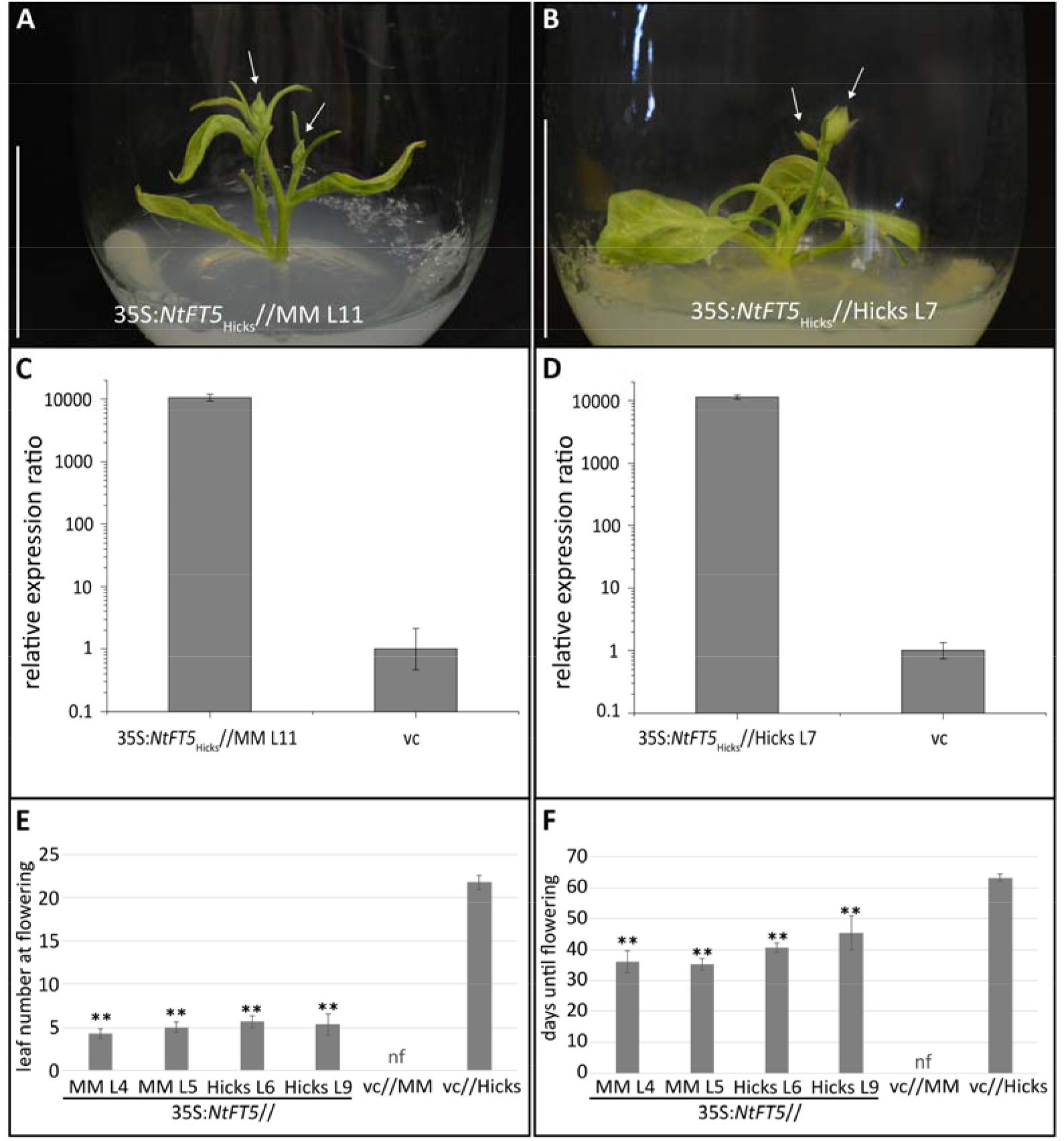
Overexpression of *NtFT5*_*Hicks*_ leads to early flowering in tobacco cultivars Maryland Mammoth (MM) and Hicks. (A,B) Constitutive overexpression of *NtFT5*_Hicks_ causes early flowering in tobacco (A) cv. MM and (B) cv. Hicks. Pictures were taken in tissue culture. Arrows indicate the formation of buds. Scale bars = 5 cm. (C,D) *NtFT5*_Hicks_ expression was analyzed by qPCR using samples isolated from entire plants harvested at the tissue culture stage. Values were normalized to the reference gene *NtEF-1α*. Expression in the vector control (vc) was set to 1 and error bars represent the combined standard deviation of target and reference gene technical replicates. *NtFT5*_Hicks_ mRNA levels were >10^4^-fold higher in MM L11 and Hicks L7 transgenic plants than the corresponding vector controls. (E) In T_1_ generation, all plants were grown in the greenhouse under LD conditions. The total number of leaves was determined when the first flower had opened. All lines of plants overexpressing *NtFT5* produced fewer leaves than the Hicks vector control. (F)The number of days until flowering was defined as the period between seed sowing and the day the first flower opened. Strong overexpression of *NtFT5* in MM and Hicks plants resulted in profound early flowering phenotypes compared to the Hicks and the non-flowering (nf) MM vector controls under LD conditions. (E,F) Mean values are shown for two 35S:*NtFT5*//MM lines (L4 and L5, n = 10), two 35S:*NtFT5*//Hicks lines (L6 n = 10, and L9, n = 9) and one vector control line for MM and Hicks (n = 10), respectively, with error bars representing ± 95% confidence intervals. Normal distribution was confirmed by applying the Kolmogorov-Smirnov test. Statistical significance was determined using a pairwise Welch’s *t*-test with Bonferroni-Holm correction (\*\**P* < 0.01). nf = non-flowering

### Functional characterization of NtFT5_MM_

The experiments described above showed that *NtFT5* expression is not suppressed in cultivar MM but that there appears to be a functional difference between the *NtFT5* genes in each cultivar. We therefore compared the *NtFT5*_Hicks_ and *NtFT5*_MM_ coding sequences. This revealed that the *NtFT5*_Hicks_ coding sequence was identical to that previously reported for cultivar SR1 (Beinecke *et al*., 2018). We therefore use the generic designation *NtFT5* hereafter when referring to the *NtFT5*_Hicks_ gene and corresponding protein. In contrast, the *NtFT5*_MM_ sequence featured a 2-bp deletion 29 bp upstream of the stop codon. The resulting frameshift mutation altered the remainder of the amino acid sequence and introduced a premature stop codon, shortening the protein sequence by six amino acids (Fig. 3A).

**Fig. 3.**
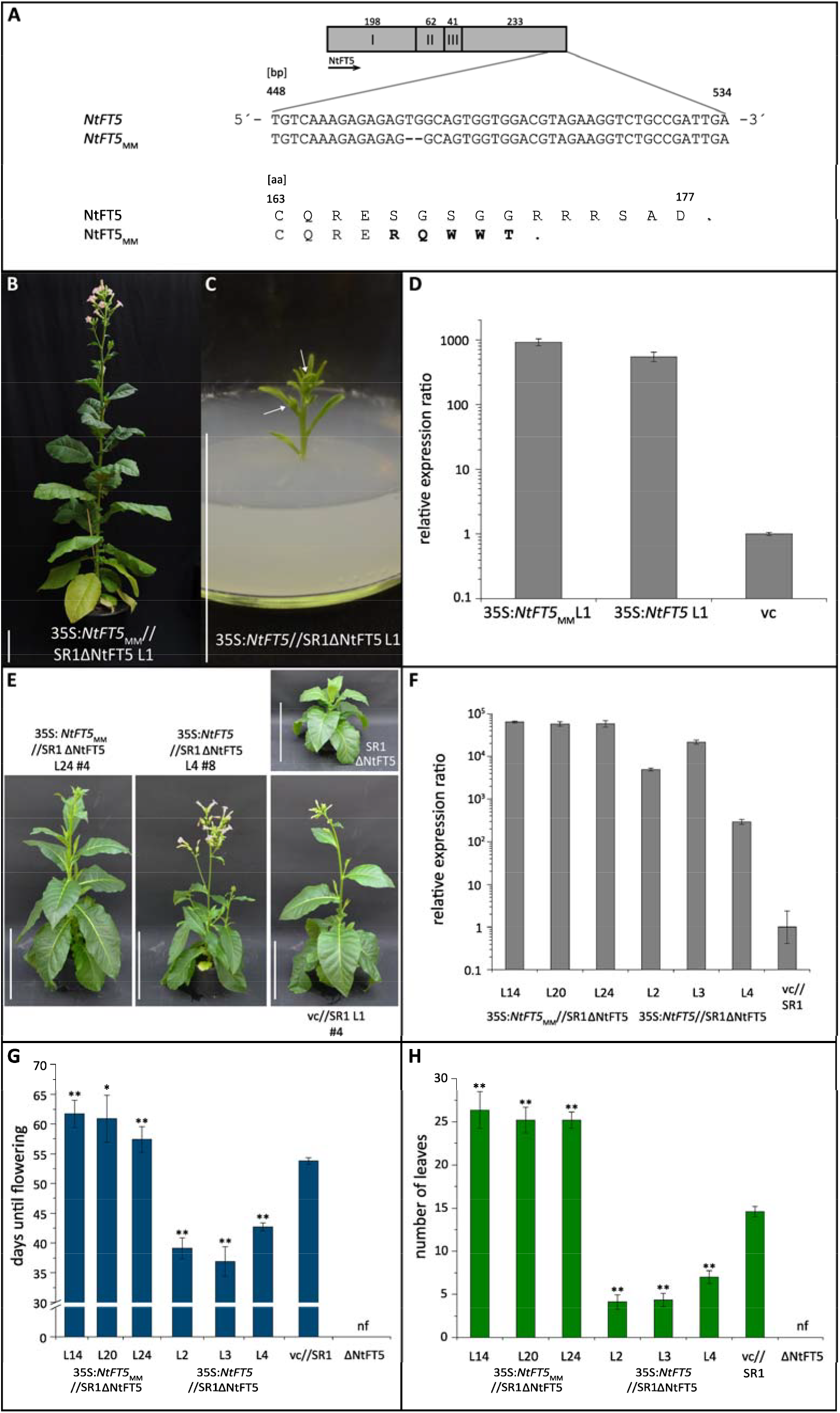
Phenotypic and molecular analysis of *NtFT5* and *NtFT5*_MM_ overexpression in SR1Δ*NtFT5* plants. (A) Exon structure of *NtFT5* and sequence alignment of the 3⍰ end of the coding sequence showing a 2-bp deletion 29 bp upstream of the typical stop codon in *NtFT5*_MM_ resulting in a truncated amino acid sequence. (B) Constitutive overexpression of *NtFT5*_MM_ in SR1Δ*NtFT5* plants led to flower development in the greenhouse when the plants reached approximately 1.2 m in height. Scale bar = 10 cm. (C) In contrast, *NtFT5* overexpression in SR1Δ*NtFT5* plants caused a profound early flowering phenotype in tissue culture. Arrows indicate the formation of buds. Scale bar = 10 cm. (D) *NtFT5*_MM_ and *NtFT5* expression was analyzed by qPCR using samples isolated from leaf material (*NtFT5*_MM_) and the whole plant shoot for *NtFT5* and vector control (vc), both harvested in tissue culture. The values were normalized to the reference gene *NtEF-1α*. Expression in the vector control was set to 1 and error bars represent the combined standard deviation of the target and reference gene technical replicates. *NtFT5*_MM_ mRNA (920-fold higher than vector control) and *NtFT5* mRNA (540-fold higher than vector control) levels strongly increased in both transgenic lines. (E) Constitutive overexpression of *NtFT5*_MM_ in the non-flowering SR1Δ*NtFT5* background caused delayed flowering compared to the wild-type SR1 vector control (vc) under LD conditions (T_1_ generation), indicating only partial complementation, whereas overexpression of *NtFT5* induced early flowering in transgenic plants. Scale bars = 30 cm. (F) *NtFT5*_MM_ and *NtFT5* expression was analyzed by qPCR using samples isolated from five pooled seedlings. The values were normalized to the reference gene *NtEF-1α*. Expression in the vector control was set to 1 and error bars represent the combined standard deviation of the target and reference gene technical replicates. The levels of *NtFT5*_MM_ mRNA were higher in all lines compared to the vector control, ranging from 58,000-fold (L20 and L24) to 64,000-fold (L14). *NtFT5* transcript levels ranged from 300-fold (L4) to 22,000-fold (L3) higher than the vector control. (G) The number of days until flowering was defined as the period between seed sowing and the day the first flower opened. Strong overexpression of *NtFT5*_MM_ in SR1Δ*NtFT5* plants resulted in delayed flowering in all transgenic lines compared to the SR1 vector control. Plants overexpressing *NtFT5* showed profound early flowering phenotypes compared the SR1 vector control. (H) The total number of leaves was determined when the first flower had opened. All lines of SR1Δ*NtFT5* plants overexpressing *NtFT5*_MM_ produced more leaves and all lines overexpressing *NtFT5* produced fewer leaves than the SR1 vector control. (G,H) Mean values are shown for three 35S:*NtFT5*_MM_//SR1Δ*NtFT5* lines (L14 and L24, n = 10; L20, n = 8), three 35S:*NtFT5*_MM_//SR1Δ*NtFT5* lines (L2 and L3, n = 9; L4, n = 10) and one vector control line (n = 10) with error bars representing ± 95% confidence intervals. Normal distribution was confirmed by applying the Kolmogorov-Smirnov test. Statistical significance was determined using a pairwise Welch’s *t*-test with Bonferroni-Holm correction (\*\**P* < 0.01; \**P* < 0.05). nf = non-flowering

We next attempted to confirm the ability of NtFT5_MM_ to act as a floral promoter. We previously generated *NtFT5*-deficient SR1 plants (SR1ΔNtFT5) using the CRISPR/Cas9 system and they were unable to flower under LD conditions (Schmidt *et al*., 2020). We therefore transformed these SR1Δ*NtFT5* plants with constructs expressing *NtFT5* or *NtFT5*_MM_ under the control of the constitutive CaMV 35S promoter (Fig. 3B–D). Ten out of 13 35S:NtFT5 transgenic plants showed a profound early flowering phenotype that failed to produce roots and thus could not be transferred to the greenhouse for the generation of seeds. In contrast, none of the 12 35S:*NtFT5*_MM_ transgenic plants flowered in tissue culture but when transferred to the greenhouse after root induction they started flowering at a later developmental stage (Fig. 3B). We analyzed transgene expression in young leaf tissue by qPCR (the primers amplified the same 190-bp fragment in the coding sequence of *NtFT5* and *NtFT5*_MM_) and detected higher NtFT5_MM_ and NtFT5 mRNA levels in the transgenic plants than in the SR1 vector control (Fig. 3D). Floral induction by *NtFT5*_MM_ was investigated in more detail by studying these transgenic N. tabacum cv. SR1ΔNtFT5 lines in the T_1_ generation (Fig. 3E–H). We cultivated three independent transgenic lines each (n = 8–10) and one SR1 vector control line (n = 10) in the greenhouse under LD conditions and documented the late and early flowering phenotype for one representative transgenic line each in comparison to a vector control (Fig. 3E). Although *NtFT5*_MM_ mRNA levels strongly increased compared to transgenic plants overexpressing *NtFT5* and the vector control (Fig. 3F), *NtFT5*_MM_ overexpression did not induce early flowering in the SR1Δ*NtFT5* plants as already observed with primary transformants. Moreover, flowering was significantly delayed in all three 35S:*NtFT5*_MM_//SR1Δ*NtFT5* lines and the number of leaves increased significantly compared to the SR1 vector control (Fig. 3G,H). In comparison, all individuals of the three transgenic lines overexpressing *NtFT5* showed a profound early flowering phenotype, resulting in significantly fewer leaves and significantly fewer days until flowering compared to the SR1 vector control with endogenous NtFT5 expression (Fig. 3G,H). This indicates that *NtFT5*_MM_ only partially complements the non-flowering phenotype of SR1Δ*NtFT5* plants, showing that *NtFT5*_MM_ is a weaker floral inducer than NtFT5.

We also transformed SR1Δ*NtFT5* and MM plants with constructs expressing *NtFT5* or *NtFT5*_MM_ under the control of the endogenous NtFT5 promoter (∼2.6kb; Supplementary Fig. S2A,B). None of these primary P-NtFT5_2.6kb_:*NtFT5* and P-NtFT5_2.6kb_:*NtFT5*_MM_ transgenic plants flowered in tissue culture or after transfer to the greenhouse (LD conditions). To overcome the non-flowering phenotype and facilitate seed production, these plants were grafted onto florally induced wild-type SR1 rootstocks. Following successful germination on selective media, we cultivated at least two independent transgenic lines per construct (n=10), one MM vector control line and MM wild-type plants (n = 5) in the greenhouse under LD conditions and documented, if the non-flowering phenotype of SR1Δ*NtFT5* and MM transgenic plants is complemented by these constructs. In the SR1Δ*NtFT5* background, 9–10 transgenic plants per line expressing P-NtFT5_2.6kb_:*NtFT5* started flowering while none of the transgenic plants expressing P-NtFT5_2.6kb_:NtFT5_MM_ in SR1Δ*NtFT5* background starting flowering (Supplementary Fig. S2A,C). When expressing *NtFT5* under the control of the endogenous NtFT5 promoter in the MM background, plants from four out of six analyzed lines started flowering in the T_1_-generation. Here, out of 10 transgenic plants, three (L5), four (L1, L8) and seven (L7) flowered within the experimental time frame (Supplementary Fig. S2B,C). In contrast, when expressing *NtFT5*_MM_ under the control of the endogenous NtFT5 promoter in the MM background, none of the seventy analyzed plants (10 plants for each of the seven analyzed lines) flowered (Supplementary Fig. S2B,C).

Thus, only expression of *NtFT5* but not *NtFT5*_MM_ under control of its own promoter could partially complement the non-flowering phenotype of SR1Δ*NtFT5* and MM plants, showing again that NtFT5_MM_ is a weaker floral inducer than NtFT5. However, as not all P-NtFT5_2.6kb_:*NtFT5*//MM plants flowered the expression of NtFT5 under control of the 2.6kb endogenous promoter fragment does not fully resemble the native NtFT5 expression.

We further examined if the homozygous, 2-bp deletion detected in *NtFT5*_MM_ in the cultivar MM correlates with a non-flowering phenotype after introgression of a functional NtFT5 allele by crossing the cultivar MM with the cultivar Hicks (Fig. 4A). We therefore grew plants of both cultivars under SD conditions to induce flowering and performed reciprocal crossings of the two cultivars. F_1_-seeds for each reciprocal cross were sown in soil and ten plants each were grown in the greenhouse under LD conditions. All plants flowered and we genotyped the plants by fragment length analysis (FLA) thereby detecting one *NtFT5* and one *NtFT5*_MM_ allele from each parental cultivar as expected. We then pooled the seeds of 10 F_1_ plants for each reciprocal cross and cultivated 95 F_2_ plants each together with the parental cultivars (n=9-10) in the greenhouse under LD. We documented the days until flowering and leaf number when the first flower had opened and genotyped each plant for the presence of the NtFT5 and NtFT5_MM_ alleles (Fig. 4 B,C; Supplementary Table S2). All plants of cultivar Hicks started flowering after 46 days with a mean number of 20 leaves, while none of the MM plants started flowering until the end of the experiment after 120 days. In this timeframe, we observed 155 flowering and 35 non-flowering F_2_ plants. Genotyping the F_2_ plants revealed that all non-flowering F_2_ plants carried the NtFT5_MM_ allele, 112 flowering F_2_ plants had one allele of each parental cultivar (NtFT5/NtFT5_MM_) and 43 flowering F_2_ plants carried only the NtFT5 allele. Interestingly, F_2_ plants with two functional NtFT5 alleles flowered on average after 52 days with a mean number of 22.5 leaves, while F_2_ plants with heterozygous NtFT5/NtFT5_MM_ alleles flowered on average after 59 days with a mean number of 26 leaves (Fig. 4 B,C). We could therefore show that the non-flowering MM phenotype under LD clearly segregates with the NtFT5_MM_ allele and that the presence of two functional NtFT5 alleles mediates earlier flowering in comparison to plants with only one functional NtFT5 allele.

**Fig. 4.**
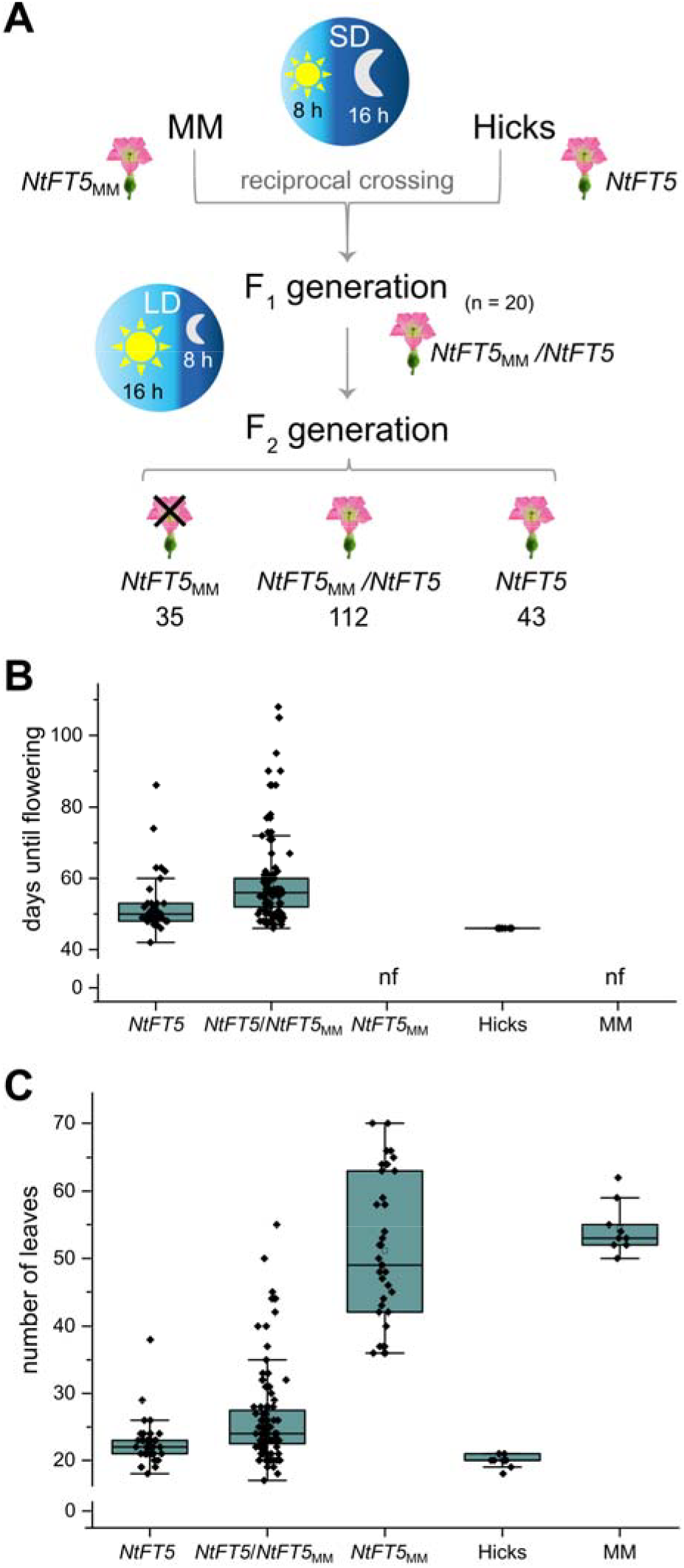
Reciprocal crossings of the cultivars Hicks and MM. (A) Experimental workflow of the crossing experiment. Flowering MM and Hicks plants were crossed reciprocally. In the F_1_ generation (n = 20) all plants flowered and showed a heterozygous NtFT5/NtFT5_MM_ genotype. Detailed phenotyping of non-flowering homozygous NtFT5_MM_ plants and flowering NtFT5 and NtFT5/NtFT5_MM_ plants was finally carried out in the F_2_ generation. (B) The number of days until flowering was defined as the period between seed sowing and the day the first flower opened. The presence of two functional NtFT5 alleles in F_2_ individuals mediated a normal flowering phenotype, while the presence of NtFT5/NtFT5_MM_ alleles mediated a delayed flowering phenotype. All F_2_ individuals with two NtFT5_MM_ alleles did not flower like the MM plants (nf = non-flowering). (C) The total number of leaves was determined when the first flower had opened (in case of non-flowering NtFT5_MM_ and MM plants at the end of the experiment after 120 days). (B,C) In the boxplots, center line = median, square = mean, box limits = upper and lower quartiles, whiskers = 1.5× interquartile range, and diamonds = individual data points.

To gain more insight into the mechanistic difference between NtFT5 and NtFT5_MM_, we determined whether the truncated NtFT5_MM_ protein was able to interact with the recently identified tobacco FD proteins (Beinecke *et al*., 2018). To initiate floral development, FT interacts with FD in the SAM (Kardailsky *et al*., 1999; Kobayashi *et al*., 1999; Abe *et al*., 2005; Wigge *et al*., 2005) and similar FT/FD interactions have been described in tobacco (Beinecke *et al*., 2018). We conducted BiFC assays in which the C-terminal portion of the fluorescent reporter protein mRFP was fused to the N-terminus of NtFD1, NtFD3, NtFD4, or NtFD1 with a mutated STAPF motif (negative control NtFD1_SAAPF_), which does not interact with tobacco FTs due to a missing phosphorylation site (Beinecke *et al*., 2018). The N-terminal portion of mRFP was fused to the N-terminus of NtFT5_MM_, NtFT5, the well-characterized AtFT as a positive control, and a truncated version of NtFT5 lacking 15 amino acids (NtFT5_FN_) as a negative control (Supplementary Method S1). The CmRFP-FD proteins were coexpressed systematically with the NmRFP-FT proteins. As expected, CLSM revealed a nuclear fluorescent signal for AtFT with any of the three native NtFD proteins (Supplementary Fig. S3) but not when NtFT5_FN*_ was combined with these native NtFDs (Fig. 5A, Supplementary Fig. S3). Furthermore, nuclear fluorescence was detected when either NtFT5_MM_ or NtFT5 was present with any of the three native NtFD proteins (NtFD1 is shown as a representative example in Fig. 5A, for the others see Supplementary Fig. S3), but not with NtFD1_SAAPF_ (Fig. 5B). We confirmed the presence of the different mRFP-FT fusion proteins by western blot analysis (Supplementary Fig. S4).

**Fig. 5.**
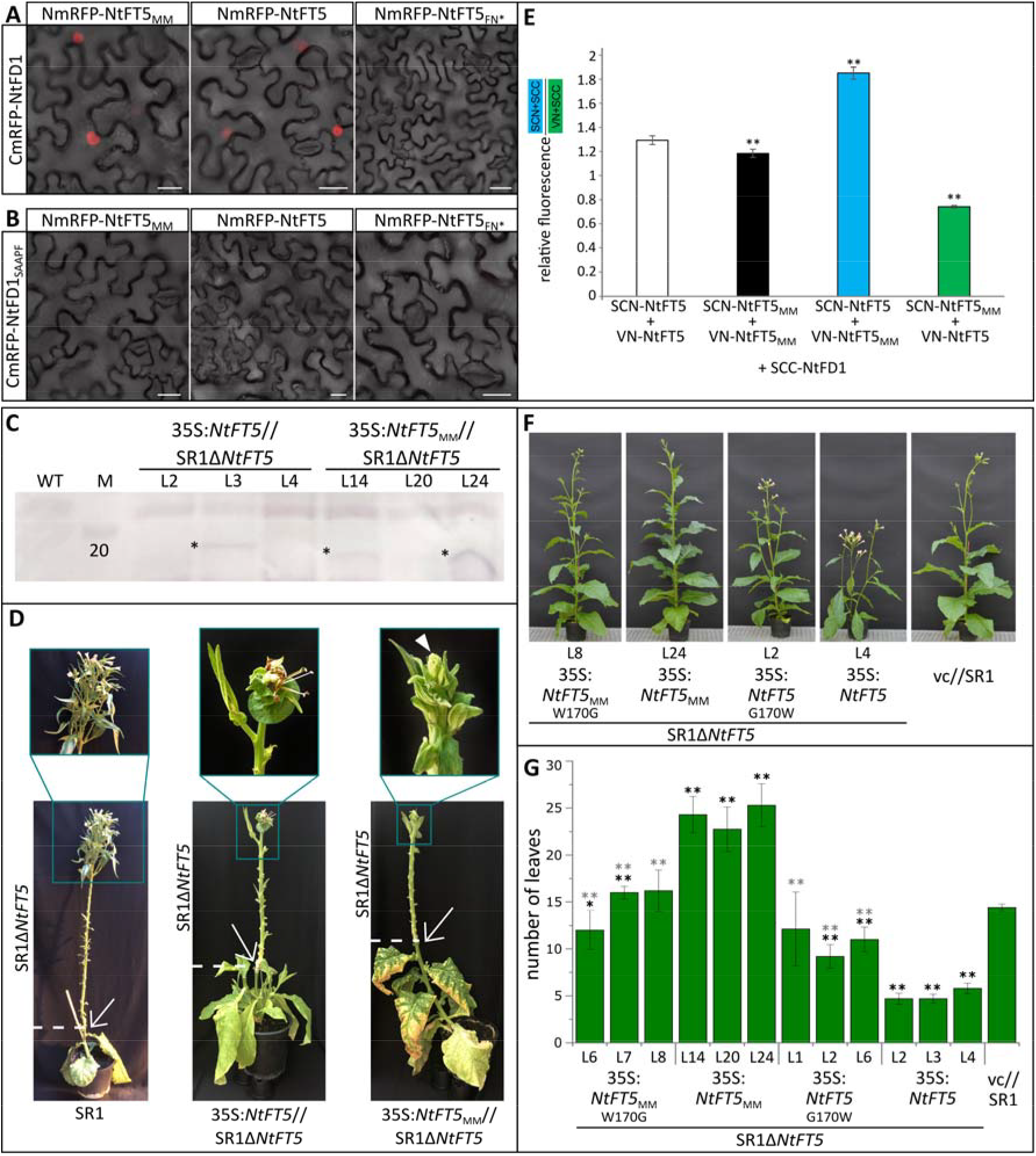
Functional characterization of NtFT5_MM_ and NtFT5. (A,B) BiFC analysis revealed that NtFD1 (A) interacts with NtFT5_MM_ and NtFT5 in *N. benthamiana* leaf epidermal cells, whereas NtFD1_SAAPF_ (B) does not interact with the tobacco FTs; 35S:*CmRFP-NtFD1* or 35S:*CmRFP-NtFD1*_SAAPF_ was co-expressed with 35S:*NmRFP* fusions of *NtFT5*_MM_, *NtFT5* and *NtFT5*_FN*_. Scale bars = 25 µm. (C) Detection of NtFT5 and NtFT5_MM_ proteins by western blotting. We loaded 50 mg of protein extracted from seedlings in each lane and detected the NtFT protein (*) with an AtFT-specific primary antibody and an alkaline phosphatase-labeled secondary antibody. The Precision Plus Protein standards were used as size markers (M). (D) Grafting experiments confirming that the NtFT5 and NtFT5_MM_ proteins are transported from wild-type (WT), 35S:*NtFT5* and 35S:*NtFT5*_MM_ stock to the SAM to induce flower-like structures on SR1Δ*NtFT5* scions. Pictures were taken when scions flowered. Emerging leaves on the scion and axillary shoots on the stock were removed regularly to enhance the source–sink gradient. (E) In mcBiFC assays, NtFD1 preferentially associated with NtFT5 rather than NtFT5_MM_. T-DNAs carrying two CaMV 35S cassettes encoding combinations of SCN and VN fusion proteins were introduced into *N. benthamiana* leaf epidermal cells together with a separate T-DNA containing a 35S:*SCC-NtFD1* construct. SCN+SCC and VN+SCC fluorescence was measured in the nuclei. Values represent the SCN+SCC signal divided by the VN+SCC signal. Error bars indicate 95% confidence intervals for means. Sample sizes (left to right): 428, 341, 535, 542. SCN, N-terminal part of S(CFP)3A; VN, N-terminal part of Venus; SCC, C-terminal part of S(CFP)3A. Statistical significance was determined using a pairwise Welch’s t-test with Bonferroni-Holm correction (\*\**P* < 0.01). (F) Replacing the tryptophan residue in NtFT5_MM_ with glycine in transgenic 35S:*NtFT5*_MM W170G_//SR1Δ*NtFT5* lines resulted in a WT-like flowering phenotype, whereas mutating the glycine residue to tryptophan in transgenic 35S:*NtFT5*_G170W_//SR1Δ*NtFT5* lines abolished the profound early flowering phenotype under LD conditions (T_1_ generation). One line each is shown as an example. (G) The number of leaves was determined when the first flower opened. All SR1Δ*NtFT5* lines overexpressing *NtFT5*_MM_ produced more leaves and all lines overexpressing *NtFT5* produced fewer leaves than the SR1 vector control. The 35S:*NtFT5*_G170W_//SR1Δ*NtFT5* lines produced more leaves and the 35S:*NtFT5*_MM G170W_//SR1Δ*NtFT5* lines produced fewer leaves than the corresponding 35S:*NtFT5*_(MM)_//SR1Δ*NtFT5* lines. Mean values are shown for three 35S:*NtFT5*_MM_//SR1Δ*NtFT5* lines (L14 and L24, n = 10; L20, n = 8), three 35S:*NtFT5*_MM_//SR1Δ*NtFT5* lines (L2, L3 and L4, n = 10), three 35S:*NtFT5*_MM W170G_//SR1Δ*NtFT5* lines (L6, L7 and L8, n = 10), three 35S:*NtFT5*_G170W_//SR1Δ*NtFT5* lines (L1, n = 8; L2 n = 10; L6, n = 6) and one vector control line (n = 10) with error bars representing ± 95% confidence intervals. Normal distribution was confirmed by applying the Kolmogorov-Smirnov test. Statistical significance was determined using a pairwise Welch’s *t*-test with Bonferroni-Holm correction (\*\**P* < 0.01; \**P* < 0.05; black = significant compared to vc; gray = significant compared to corresponding native construct).

Next, we determined whether the C-terminal truncation of NtFT5_MM_ affects protein stability. Crude protein extracts of T_1_ seedlings from three independent transgenic lines (same lines as in Fig. 3) and SR1 wild-type seedlings as controls were used for western blotting. We detected NtFT5 and NtFT5_MM_ using the AtFT-specific antibody only in crude protein extracts of transgenic seedlings overexpressing the corresponding NtFT. Specific signals were present for one 35S:*NtFT5*//SR1Δ*NtFT5* line (L3), as well as two 35S:*NtFT5*_MM_//SR1Δ*NtFT5* lines (L14 and L24, Fig. 5C). The band for *NtFT5*_MM_ samples migrated further, reflecting its lower molecular weight due to the truncated C-terminus. This experiment demonstrated that the truncated C-terminus of *NtFT5*_MM_ does not appear to affect protein stability.

Next, we investigated whether the truncation of NtFT5_MM_ impairs the transport of this protein via the phloem. We therefore grafted SR1Δ*NtFT5* scions onto rootstocks of either 35S:NtFT5//SR1Δ*NtFT5* or 35S:*NtFT5*_MM_//SR1Δ*NtFT5* lines as well as wild-type control plants (n = 3 each). All SR1Δ*NtFT5* scions grafted onto wild-type control stocks flowered 7–9 weeks after grafting, with 35–49 leaves/nodes produced on the scion (Fig. 5D). After approximately 11 weeks, one SR1Δ*NtFT5* scion grafted onto 35S:*NtFT5*//SR1Δ*NtFT5* stock, and two grafted onto 35S:*NtFT5*_MM_//SR1Δ*NtFT5* stocks, showed floral-like structures, with 21 and 24/25 leaves/nodes produced on the scion, respectively (Fig. 5D). However, only 50% of these grafts showed floral transition of the SAM, and complete inflorescences (as induced by wild-type stocks) were not formed. Given that 35S:*NtFT5*//SR1Δ*NtFT5* stocks produced many axillary shoots (which we removed regularly) this may have impaired source–sink transport to the scion. For the 35S:*NtFT5*_MM_//SR1Δ*NtFT5* stocks, this might indicate a functionally impaired NtFT5_MM_ protein. These results suggest that NtFT5_MM_ is transported to the shoot apex like NtFT5 but floral transition mediated by 35S:*NtFT5*_MM_//SR1Δ*NtFT5* stocks was slightly delayed compared to floral transition of the SR1Δ*NtFT5* scion grafted onto a 35S:*NtFT5*//SR1Δ*NtFT5* stock, again matching our earlier results that NtFT5_MM_ is a weaker floral inducer.

Because the transport of NtFT5_MM_ was apparently normal, we conducted mcBiFC assays to test the binding affinity of both NtFTs with NtFD1. Using this assay, we recently confirmed the lower binding affinity of floral repressor NtFT2 to NtFD1 compared to floral activator NtFT4 (Beinecke *et al*., 2018). We therefore prepared two cassettes controlled by the CaMV 35S promoter for the expression of NtFT5 or NtFT5_MM_ as N-terminal fusions with the N-terminal component of cyan fluorescent protein S(CFP)3A (SCN) together with an N-terminal fusion of NtFT5 or NtFT5_MM_ with the N-terminal component of Venus (VN) (Supplementary Fig. S5). A construct in which the N-terminal portion of NtFD1 was fused to the C-terminal portion of S(CFP)3A (SCC) was carried by a separate bacterial strain. Interactions between SCN and SCC reconstitute S(CFP)3A, which emits blue fluorescence when excited, whereas chimeric association between VN and SCC emits green fluorescence. After agroinfiltration and the cultivation of *N. benthamiana* under constant light for 3 days, abaxial epidermal leaf cells were analyzed by CLSM and the nuclear signal intensity was measured. The co-expression of SCN-NtFT5 and VN-NtFT5 with SCC-NtFD1, as well as SCN-NtFT5_MM_ and VN-NtFT5_MM_ with SCC-NtFD1, resulted in little difference between the blue and green signals and served as an equilibrium control with balanced competition (Fig. 5E and Supplementary Fig. S5). However, when SCN-NtFT5, VN-NtFT5_MM_ and SCC-NtFD1 were co-expressed, a significant shift towards the blue signal of SCN/SCC was detected, whereas the co-expression of SCN-NtFT5_MM_, VN-NtFT5 and SCC-NtFD1 produced a dominant green VN/SCC signal. This suggests that NtFT5_MM_ may have a lower binding affinity than NtFT5 for NtFD1, and that NtFD1 may preferably associate with NtFT5 because it is more competitive at the protein level than NtFT5_MM_ with its truncated C-terminus.

Our results show that the C-terminal amino acid residues of NtFT5 fulfil an important role in floral transition. Arabidopsis mutants with the substitution G171E in the corresponding region have a late flowering phenotype (Koornneef *et al*., 1991; Kardailsky *et al*., 1999; Kobayashi *et al*., 1999). In NtFT5_MM_, a tryptophan residue is present at the corresponding glycine position (G170W) so we designed overexpression constructs with reciprocal amino acid substitutions (namely *NtFT5*_G170W_ and *NtFT5*_MM W170G_) and analyzed these in transgenic SR1ΔNtFT5 lines (Supplementary Fig. S6A). We regenerated three independent 35S:NtFT5_G170W_ and 35S:*NtFT5*_MM W170G_ transgenic lines each and investigated these lines in the T_1_ generation compared to 35S:*NtFT5* and 35S:*NtFT5*_MM_ transgenic lines in more detail (Fig. 5F,G and Supplementary Fig. S6B,C). We cultivated three independent transgenic lines each (n = 6–10) and one SR1 vector control line (n = 10) in the greenhouse under LD conditions and documented the flowering phenotype for one representative transgenic line each in comparison to a vector control (Fig. 5F). The *NtFT5*_G170W_ and *NtFT5*_MM W170G_ mRNA levels were much higher than the vector control and were in the range of the *NtFT5*_MM_ mRNA overexpression levels (Supplementary Fig. S6B). However, *NtFT5*_MM W170G_//SR1Δ*NtFT5* lines flowered significantly earlier and the number of leaves was significantly lower compared to the 35S:*NtFT5*_MM_//SR1Δ*NtFT5* lines (Fig. 5G and Supplementary Fig. S6C). Accordingly, 35:*NtFT5*_G170W_//SR1Δ*NtFT5* lines flowered later than 35S:*NtFT5*//SR1Δ*NtFT5* overexpressing plants, resulting in the production of significantly more leaves and significantly more days until flowering (Fig. 5G and Supplementary Fig. S6C). This indicates that both substitutions at position 170, tryptophan to glycine (in NtFT5_MM W170G_) and glycine to tryptophan (in NtFT5_G170W_), produced intermediate-strength floral inducers stronger than NtFT5_MM_ but weaker than NtFT5.

## Discussion

Many plant species initiate the developmental switch from vegetative to reproductive growth under specific photoperiodic conditions. Although tobacco is a day-neutral species, floral initiation depends on the photoperiod-dependent expression of various *FT* genes (Harig *et al*., 2012; Beinecke *et al*., 2018). Previous studies have mostly focused on *FT* genes expressed under SD conditions, but the recently identified NtFT5 gene is expressed under both SD and LD conditions. *NtFT5* has been characterized as the major floral activator under LD conditions in tobacco cultivar SR1, making this protein an interesting candidate for further analysis in the SD cultivar MM (Beinecke *et al*., 2018; Schmidt *et al*., 2020).

Comparative expression analysis revealed that *NtFT5* is expressed at similar levels in cultivar MM and the day-neutral cultivar Hicks, which excludes the possibility that the LD-specific non-flowering phenotype of MM is caused by the silencing of this locus (Fig. 1I,J). We also found that the *NtFT5* coding sequence was identical in cultivars Hicks and SR1, and that the overexpression of this sequence in MM permitted flowering under LD conditions (Fig. 2), suggesting that the *NtFT5*_MM_ protein may be functionally impaired. Further analysis revealed a 2-bp deletion near the 3⍰ end of the *NtFT5*_MM_ gene causing a frameshift that introduced a premature termination codon and an altered C-terminus (Fig. 3A). By analyzing the F_2_ offspring of the reciprocal crossings between the cultivars Hicks and MM we showed that this 2-bp deletion in the *NtFT5*_MM_ gene is tightly linked with a non-flowering phenotype under LD if present homozygously (*NtFT5*_MM_/ *NtFT5*_MM_) and confers delayed flowering if present heterozygously (*NtFT5*_MM_/ *NtFT5*) (Fig. 4). A similar effect was also described for *NtFT5* knockout mutations (Schmidt *et al*., 2020). The 2-bp deletion near the 3⍰ end of the *NtFT5*_MM_ gene results in a *NtFT5*_MM_ protein that lacks the six amino acids normally found at the C-terminus and exhibits five amino acids substitutions at the C-terminus (Fig. 3A). Because the NtFT5_MM_ protein could be detected by western blot and floral structures at the SAM of SR1Δ*NtFT5* scions grafted onto 35S:*NtFT5*_MM_//SR1Δ*NtFT5* stocks were induced (Fig. 5C,D), we concluded that neither protein stability nor transport is affected by the C-terminal truncation and alteration of *NtFT5*_MM_. We tested the ability of NtFT5 and *NtFT5*_MM_ to induce flowering under LD conditions by expressing each variant in SR1Δ*NtFT5* plants. As expected, NtFT5 overexpression resulted in a profound early flowering phenotype (Fig. 3B,G,H), as described for transgenic tobacco lines overexpressing other floral activators such as NtFT4 and AtFT (Harig *et al*., 2012; Beinecke *et al*., 2018). In contrast, the overexpression of *NtFT5*_MM_ caused plants to flower at a later developmental stage, even though *NtFT5*_MM_ mRNA levels were higher than *NtFT5* mRNA levels in the corresponding transgenic SR1Δ*NtFT5* plants (Fig. 3C,G,H). The late-flowering behavior of transgenic 35S:*NtFT5*_MM_//SR1Δ*NtFT5* compared with the normal flowering time of SR1 vector control lines (expressing *NtFT5* at the endogenous level) showed that a fully functional endogenous *NtFT5* gene can more effectively induce flowering than the strong overexpression of *NtFT5*_MM_. In line with, this we could show that only the expression of *NtFT5* under control of its endogenous promoter could complement the non-flowering phenotypes of SR1Δ*NtFT5* and MM plants while expression of P-NtFT5_2.6kb_:*NtFT5*_MM_ did not induce flowering (Supplementary Fig. S2). This suggests that the truncated and altered C-terminus of *NtFT5*_MM_ weakens its function as a floral activator.

As previously shown in Arabidopsis and rice, FT proteins interact with FD proteins to regulate flower development (Abe *et al*., 2005; Wigge *et al*., 2005; Taoka *et al*., 2011). Similarly, the tobacco FT proteins interact with three FD proteins, and it was therefore important to determine whether the truncated NtFT5_MM_ protein retained this ability (Beinecke *et al*., 2018). Our BiFC experiments showed that both NtFT5 and NtFT5_MM_ interacted with all three tobacco FD proteins, but mcBiFC experiments revealed that NtFD1 may preferably associate with NtFT5 because it is more competitive than NtFT5_MM_ at the protein level (Fig. 5A,B,E; Supplementary Fig. S3). It is widely accepted that 14-3-3 proteins mediate interactions between FT and FD (Pnueli *et al*., 2001; Ho and Weigel, 2014; Taoka *et al*., 2011; Collani *et al*., 2019; Zuo *et al*., 2021) and the amino acid residues critical for this interaction are highly conserved in FT-like proteins from angiosperms and are located upstream of the 2-bp deletion in *NtFT5*_MM_. However, our results suggest that additional amino acids at the NtFT5 C-terminus are required for proper interaction with NtFD proteins. Furthermore, the C-terminal residues of NtFT5 fulfil an important role in floral activation as shown by the late flowering phenotype of AtFT mutants with the substitution G171E (Koornneef *et al*., 1991; Kardailsky *et al*., 1999; Kobayashi *et al*., 1999). In NtFT5_MM_, the corresponding glycine residue is replaced with tryptophan (G170W), suggesting that the truncated and altered C-terminus of NtFT5_MM_ may influence its regulatory activity or binding affinity for NtFD1. Indeed, replacing the tryptophan residue in NtFT5_MM_ with glycine in transgenic 35S:NtFT5_MM W170G_//SR1Δ*NtFT5* lines restored wild-type flowering behavior, whereas mutating the glycine residue to tryptophan in transgenic 35S:*NtFT5*_G170W_//SR1Δ*NtFT5* lines abolished the profound early flowering phenotype (Fig. 5F,G). Furthermore, experiments in Arabidopsis have shown that the overexpression of a truncated FT gene yielding a protein lacking the last seven amino acids (FTΔ168) causes a late flowering phenotype in *ft-10* mutant plants compared to *ft-10* plants overexpressing functional *FT* (Kim *et al*., 2016). FTΔ168 and NtFT5_MM_ both lack the glycine residue at position 170/171, which is strongly conserved in the FT-like family. Interestingly, members of the TFL1-like family of floral repressors have a highly conserved C-terminus that lacks the glycine residue at the corresponding position (Supplementary Fig. S7). However, the glycine residue is present in both floral activators and floral inhibitors of the FT-like family, so its precise function in terms of floral regulation remains unclear. Additional amino acid residues that are necessary for the regulatory activity of FT proteins have been described (Hanzawa *et al*., 2005; Ahn *et al*., 2006; Ho and Weigel, 2014) and it seems likely that the presence of all these conserved residues is required for the correct function of FT-like floral activators. However, further studies are needed to understand in detail the role of the C-terminus of FT in flower development.

MM plants can flower under SD conditions, indicating that the high levels of mRNA encoding the strong activator NtFT4 and the weak activator NtFT5_MM_ under SD conditions are sufficient to induce flowering, whereas the low levels of both transcripts under LD conditions are not (Fig. 1I,J). Only the strong overexpression of *NtFT5*_MM_ can overcome the non-flowering phenotype in SR1ΔNtFT5 plants (Fig. 3). However, even strong overexpression can only complement the non-flowering phenotype to a certain extent, given that the transgenic 35S:*NtFT5*_MM_//SR1Δ*NtFT5* plants still induce flower development at a later time point compared to the SR1 vector control. Under SD conditions, the MM plants flowered one week later than the day-neutral cultivar Hicks (Fig. 1). RNA interference (RNAi) experiments in tobacco SR1 plants have shown that the silencing of *NtFT4* or *NtFT5* can delay flowering, but the effect was stronger for *NtFT5* (Beinecke *et al*., 2018). This suggests that NtFT5 has a stronger effect than NtFT4 in flower development and explains the delayed flowering phenotype of cultivar MM compared to day-neutral cultivar Hicks under SD conditions, given that the latter still produces a functional NtFT5 protein.

Taken together, our results demonstrate that NtFT5_MM_ acts as a weak floral activator even when strongly overexpressed, which may reflect its lower binding affinity for NtFD. Thus, a possible model for the MM phenotype is that, under LD conditions, the typical levels of *NtFT5*_MM_ mRNA are insufficient for floral induction, but under SD conditions floral induction in MM plants is facilitated by the high levels of *NtFT4* mRNA, resulting in the switch from vegetative to reproductive growth. Thus, our results provide insight into the SD-specific flowering behavior of MM, a tobacco cultivar that was instrumental in the discovery of photoperiodism and was first described a century ago.

## Abbreviations

(FT): FLOWERING LOCUS T
(LD): long-day
(MM): Maryland Mammoth
(SD): short-day

## Supplementary data

**Fig. S1**. Detection of *NtFT7* in tobacco (*N. tabacum*) cultivars Maryland Mammoth (MM), Hicks and SR1, and *N. sylvestris*.

**Fig. S2**. Phenotypic analyses of SR1ΔNtFT5 and MM plants expressing *NtFT5* or *NtFT5*_MM_ under control of the endogenous promoter P-NtFT5_2.6kb_.

**Fig. S3**. BiFC analysis of NtFT5_R61G_, NtFT5_R129A_, NtFT5_FNC*_ and AtFT with NtFD1 as well as NtFD3 and NtFD4 with NtFT5_MM_, NtFT5, NtFT5_FN*_ and AtFT.

**Fig. S4**. Western blot analysis of BiFC experiments with different mRFP-FT fusion proteins and NtFD1.

**Fig. S5.** Multicolor BiFC assay.

**Figure S6**. Functional characterization of NtFT5_MM W170G_ and NtFT5_G170W_ mutants.

**Figure S7.** Amino acid sequence alignment of FT-like and TFL-like proteins.

**Table S1.** List of oligonucleotides used in this study.

**Table S2**. Geno- and phenotype of F_2_ individuals.

**Methods S1**. Cloning procedures for BiFC.

## Acknowledgements

We acknowledge the technical support of Michael Lahme, Christiane Fischer, Heike Hinte, Dewi Hadisoetanto, Sai Aparna Nagarajan, Sascha Ahrens and Andreas Wagner.

## Author contributions

LG, AK, DP, GAN: conceptualization, project administration and supervision; LG, MMZ, AK, AS, JM: investigation; DRW: formal analysis; LG, MMZ: visualization; LG, MMZ, AK, RMT, GAN: writing

## Data availability

All data supporting the findings of this study are available within the manuscript and its supplementary materials.

