## Supplement for "A mutant allele of the long-day flowering promoter NtFT5 unravels the mystery behind the short-day-specific flowering of tobacco cultivar Maryland Mammoth"

**Supplementary Information**

**Figure S1.** Detection of *NtFT7* in tobacco (*N. tabacum*) cultivars Maryland Mammoth (MM), Hicks and SR1, and *N. sylvestris*.

**Figure S2.** Expression of NtFT5 and NtFT5_MM_ under control of the endogenous promoter P-NtFT5_2.6kb_ in SR1ΔNtFT5 or MM background.

**Figure S3.** BiFC analysis of NtFT5_R61G_, NtFT5_R129A_, NtFT5_FNC*_ and AtFT with NtFD1 as well as NtFD3 and NtFD4 with NtFT5_MM_, NtFT5, NtFT5_FN*_ and AtFT.

**Figure S4.** Western blot analysis of BiFC experiments with different mRFP-FT fusion proteins and NtFD1.

**Figure S5.** Multicolor BiFC assay.

**Figure S6.** Functional characterization of NtFT5_MM W170G_ and NtFT5_G170W_ mutants.

**Figure S7.** Amino acid sequence alignment of FT-like and TFL-like proteins.

**Table S1.** List of oligonucleotides used in this study.

**Methods S1.** Cloning procedures for BiFC.

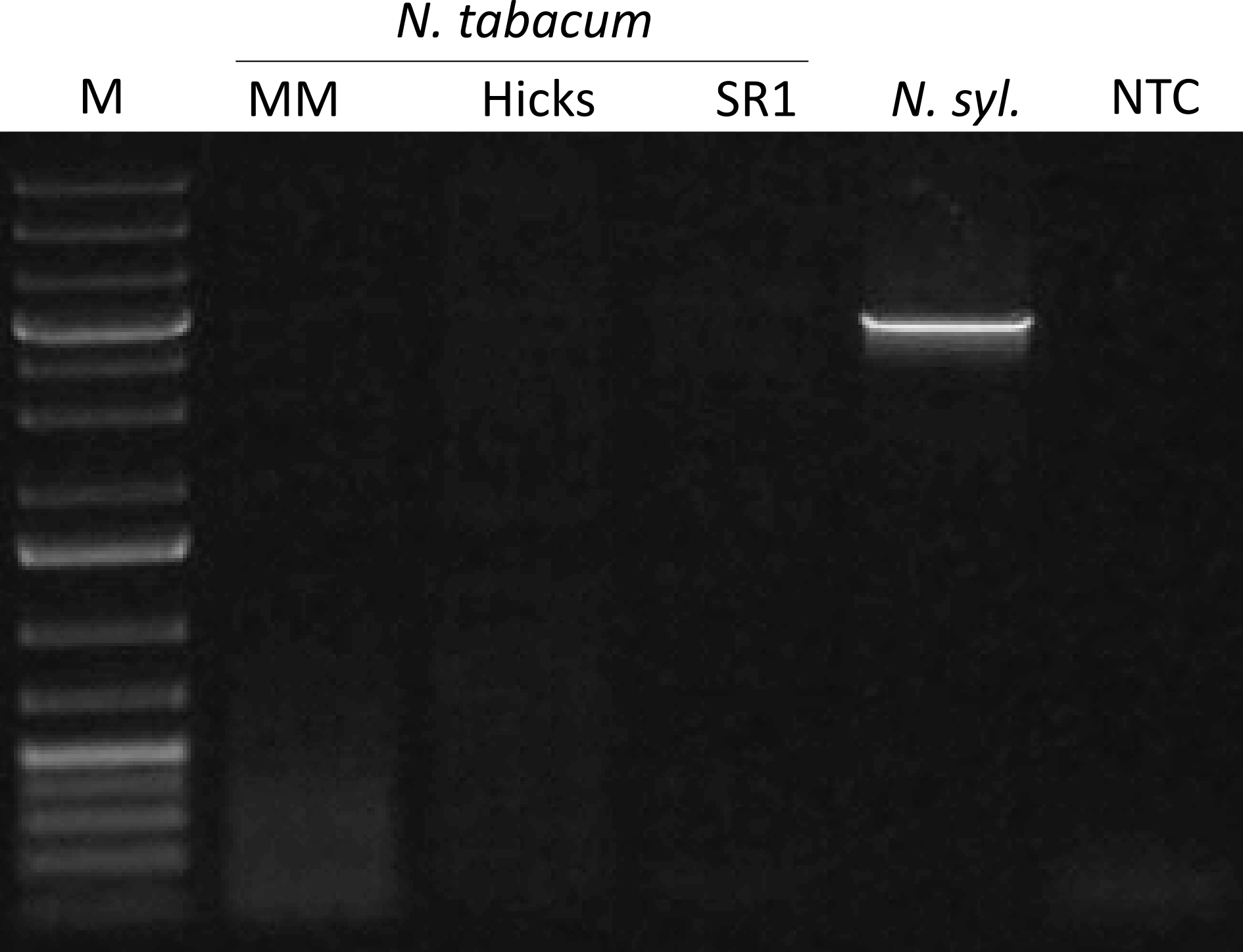

**Figure S1.** **Detection of *NtFT7* in tobacco (*N. tabacum*) cultivars Maryland Mammoth (MM), Hicks and SR1, and *N. sylvestris*.**

PCR analysis using genomic DNA revealed a band for *NtFT7* in *N. sylvestris* cv. Speg. & Comes (*N. syl.*) but not in any of the *N. tabacum* cultivars. PCR was carried out using the primers listed in Supplementary Table S1. PCR products were separated by 1% (w/v) agarose gel electrophoresis with the Gene Ruler 1 kb DNA Ladder (Thermo Fisher Scientific) as the size marker (M). NTC = no-template control.

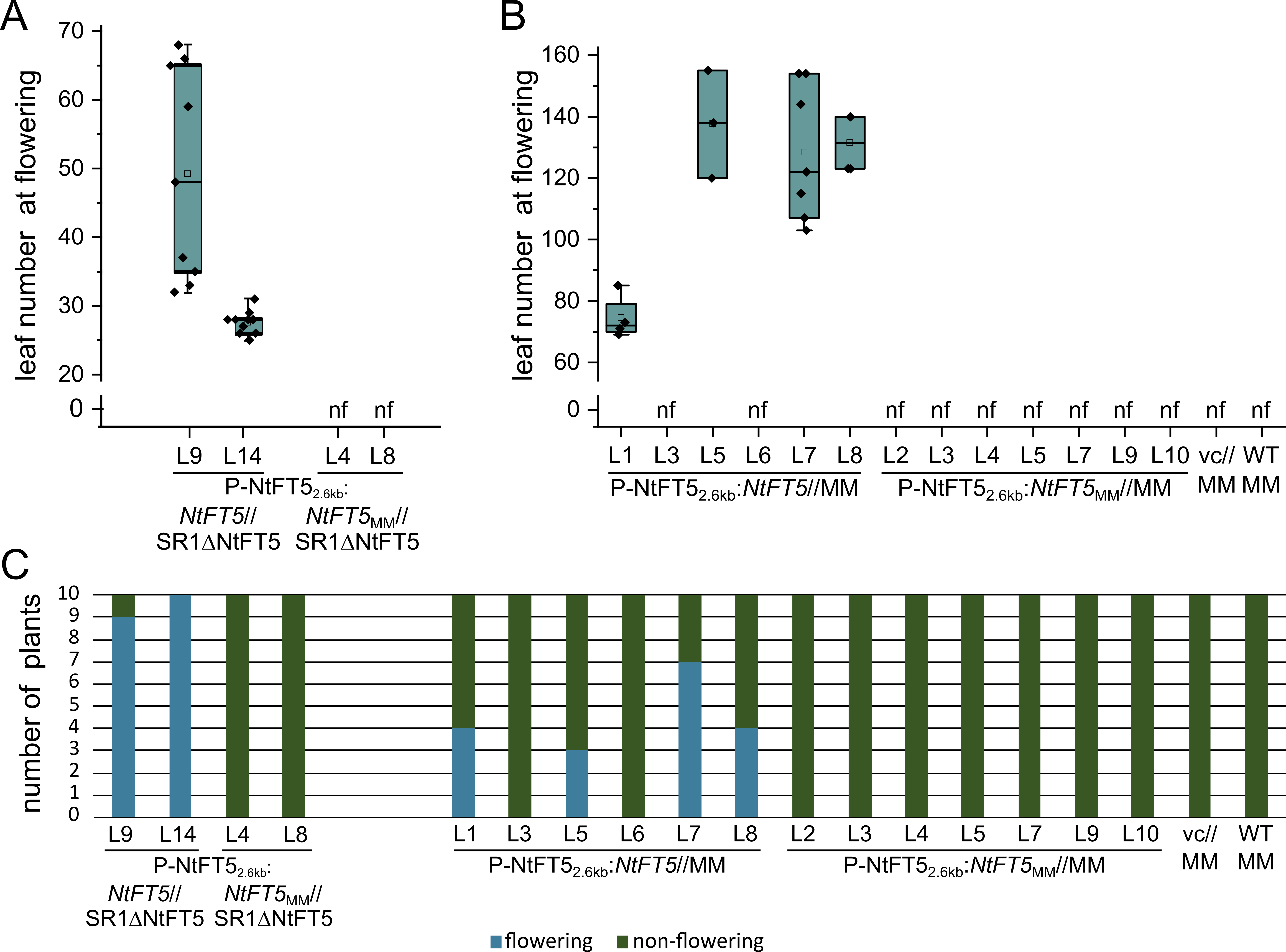

**Figure S2. Phenotypic analyses of SR1ΔNtFT5 and MM plants expressing *NtFT5* or *NtFT5*_MM_ under control of the endogenous promoter P-NtFT5_2.6kb_.** **A)** Flowering phenotype of SR1ΔNtFT5 plants expressing NtFT5 or NtFT5MM under control of a 2.6 kb fragment of the *NtFT5*-promoter. **B)** Flowering phenotype of MM plants expressing NtFT5 or NtFT5MM under control of a 2.6 kb fragment of the *NtFT5*-promoter. (A,B) Leaf number was determined when first flower opened. nf: non-flowering at the end of the experiment (xx days after transfer to the greenhouse). **C)** Percentage of flowering (blue) and non-flowering (green) plants for each line.

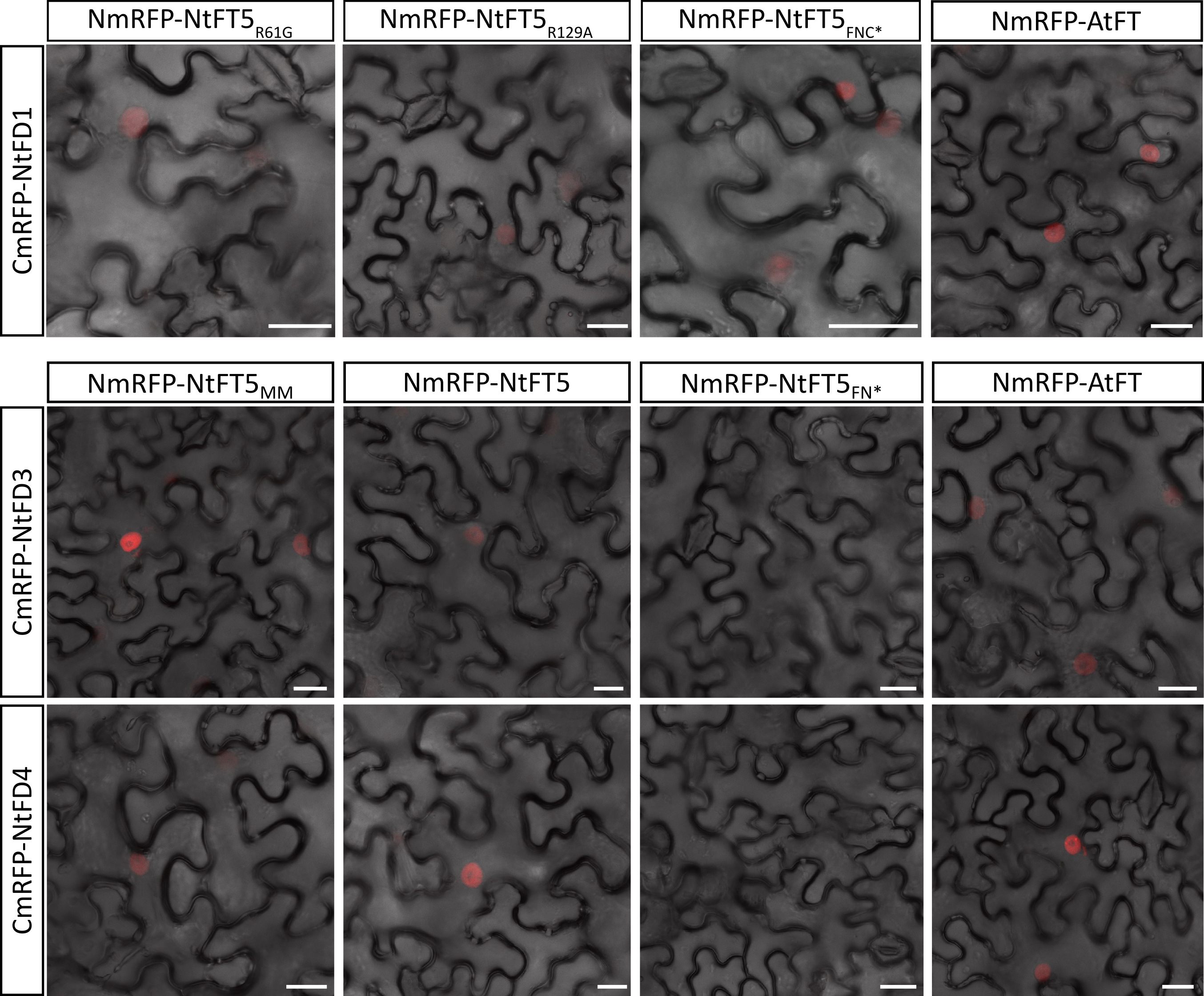

**Figure S3. BiFC analysis of NtFT5_R61G_, NtFT5_R129A_, NtFT5_FNC*_ and AtFT with NtFD1 as well as NtFD3 and NtFD4 with NtFT5_MM_, NtFT5, NtFT5_FN*_ and AtFT.**

The 35S:*CmRFP-NtFD* constructs were co-expressed with 35S:*NmRFP* fusions of *NtFT5*_R61G_, *NtFT5*_R129A_, *NtFT5*_FNC*_, *AtFT*, *NtFT5*_MM_, *NtFT5* and *NtFT5*_FN*_. BiFC analysis confirmed that NtFD1 interacts with NtFT5_R61G_, NtFT5_R129A_, NtFT5_FNC*_ and AtFT, and also that both NtFD3 and NtFD4 interact with NtFT5_MM_, NtFT5 and AtFT in *N. benthamiana* leaf epidermal cells. No interaction was detected for NtFT5_FN*_ with NtFD1, NtFD3 or NtFD4. Scale bars = 25 µm.

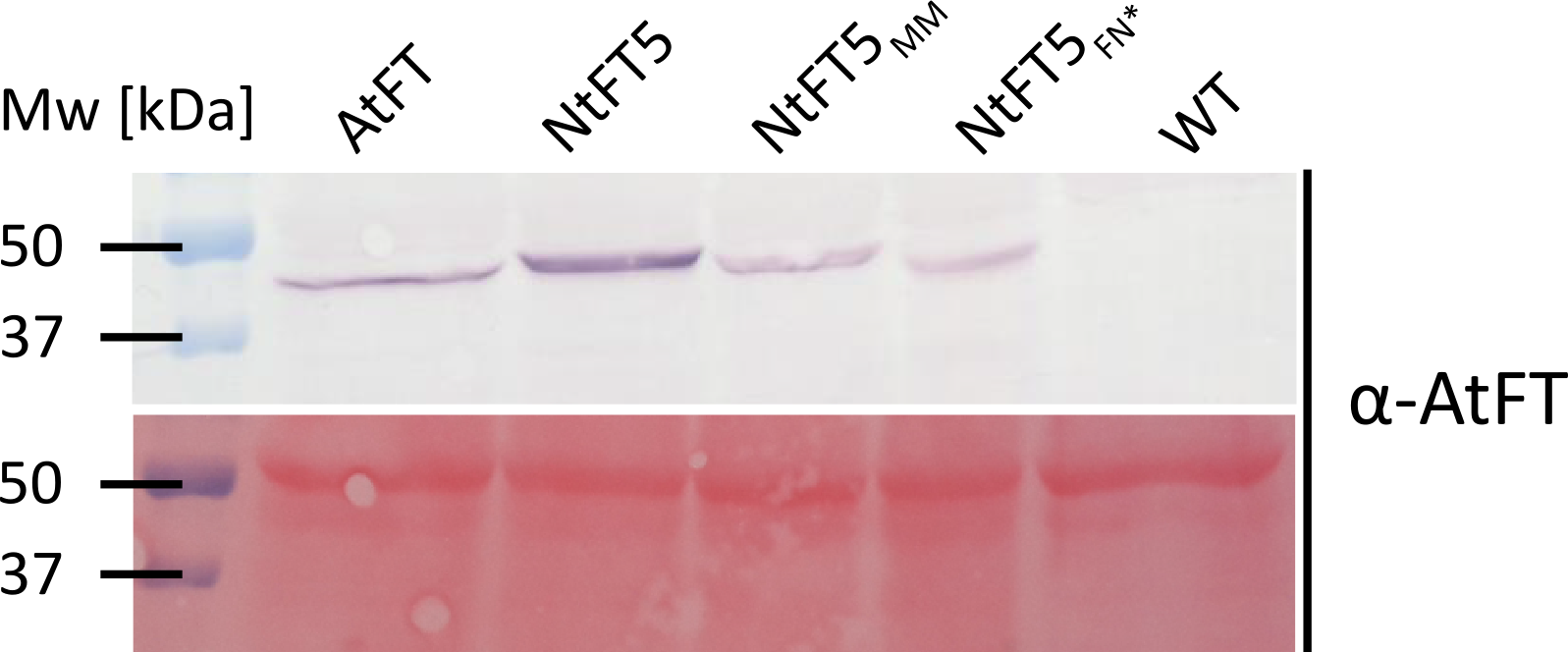

Figure S4. Western blot analysis of BiFC experiments with different mRFP-FT fusion proteins and NtFD1. We carried out western blot analysis using 12-μg protein extracts from infiltrated leaf epidermis. Using the specific primary antibody against AtFT, we detected a band at the anticipated molecular weight for mRFP fusions of AtFT, NtFT5, NtFT5_MM_ and NtFT5_FN*_ in the BiFC experiments with NtFD1, indicating that all FT proteins are stable. The bands were visualized using an alkaline phosphatase-labeled secondary antibody. Precision Plus Protein standards were used as size markers (Mw).

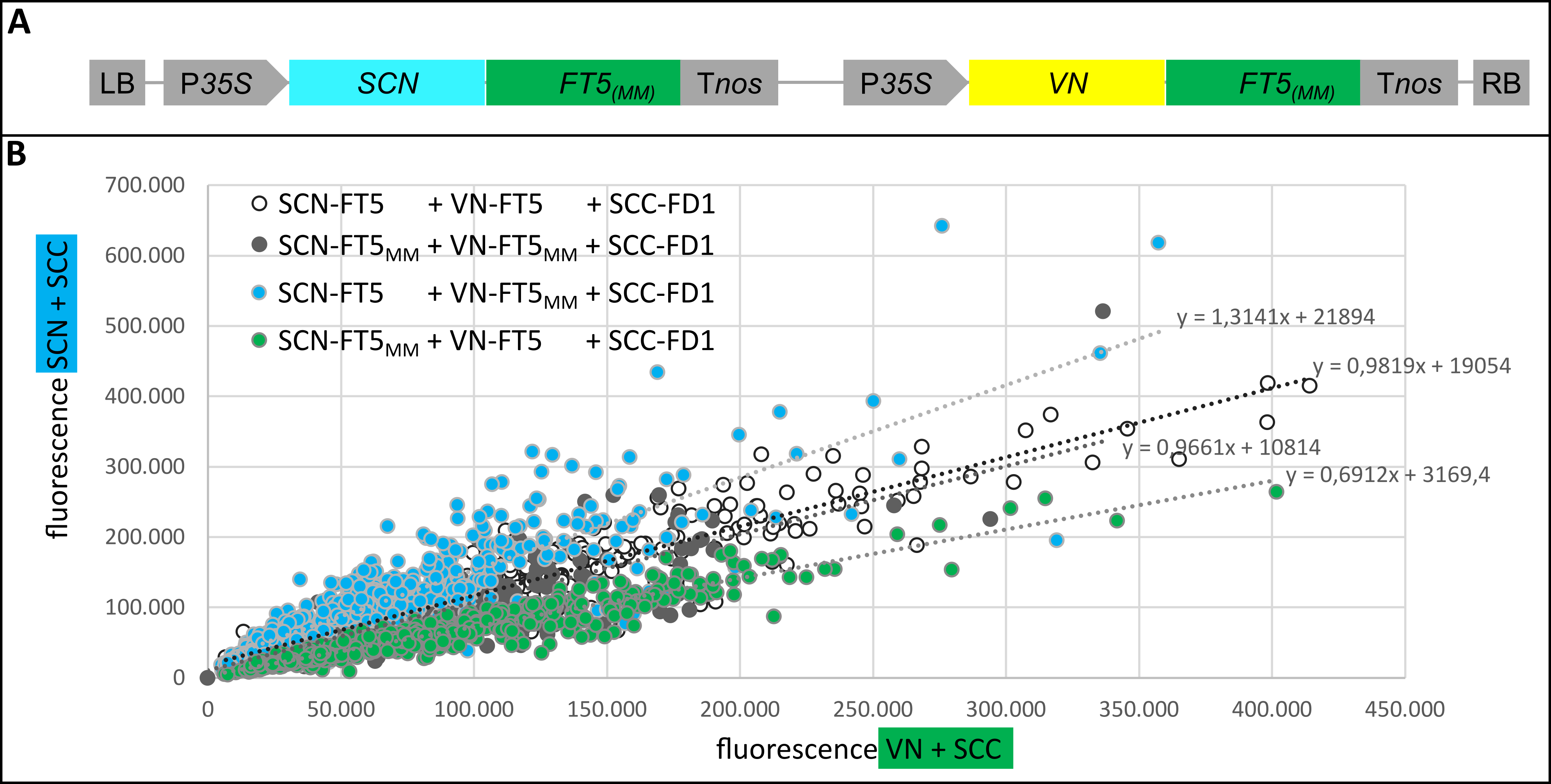

**Figure S5. Multicolor BiFC (mcBiFC) assay.** (A) Schematic map of the T-DNA carrying two expression cassettes. (B) Analysis of different combinations of NtFT5 and NtFT5_MM_ fusion proteins interacting with an SCC-NtFD1 fusion. Each dot represents a single nucleus. The *x*-axis indicates the strength of the VN+SCC signal, whereas the *y*-axis indicates the strength of the SCN+SCC signal. The combinations SCN-NtFT5 + VN-NtFT5 + SCC-NtFD1 (white; n = 428) and SCN-NtFT5_MM_ + VN-NtFT5_MM_ + SCC-NtFD1 (black; n = 341) show balanced competition. The combinations SCN-NtFT5 + VN-NtFT5_MM_ + SCC-NtFD1 (blue; n = 535) and SCN-NtFT5_MM_ + VN-NtFT5 + SCC-NtFD1 (green; n = 542) are both shifted towards the signal of the NtFT5-NtFD1 interaction.

Abbreviations: SCN = N-terminal part of S(CFP)3A; VN = N-terminal part of Venus; SCC = C-terminal part of S(CFP)3A; P*35S =* CaMV 35S promoter; T*nos =* *A. tumefaciens* *nopaline synthase* terminator; LB = left border; RB = right border.

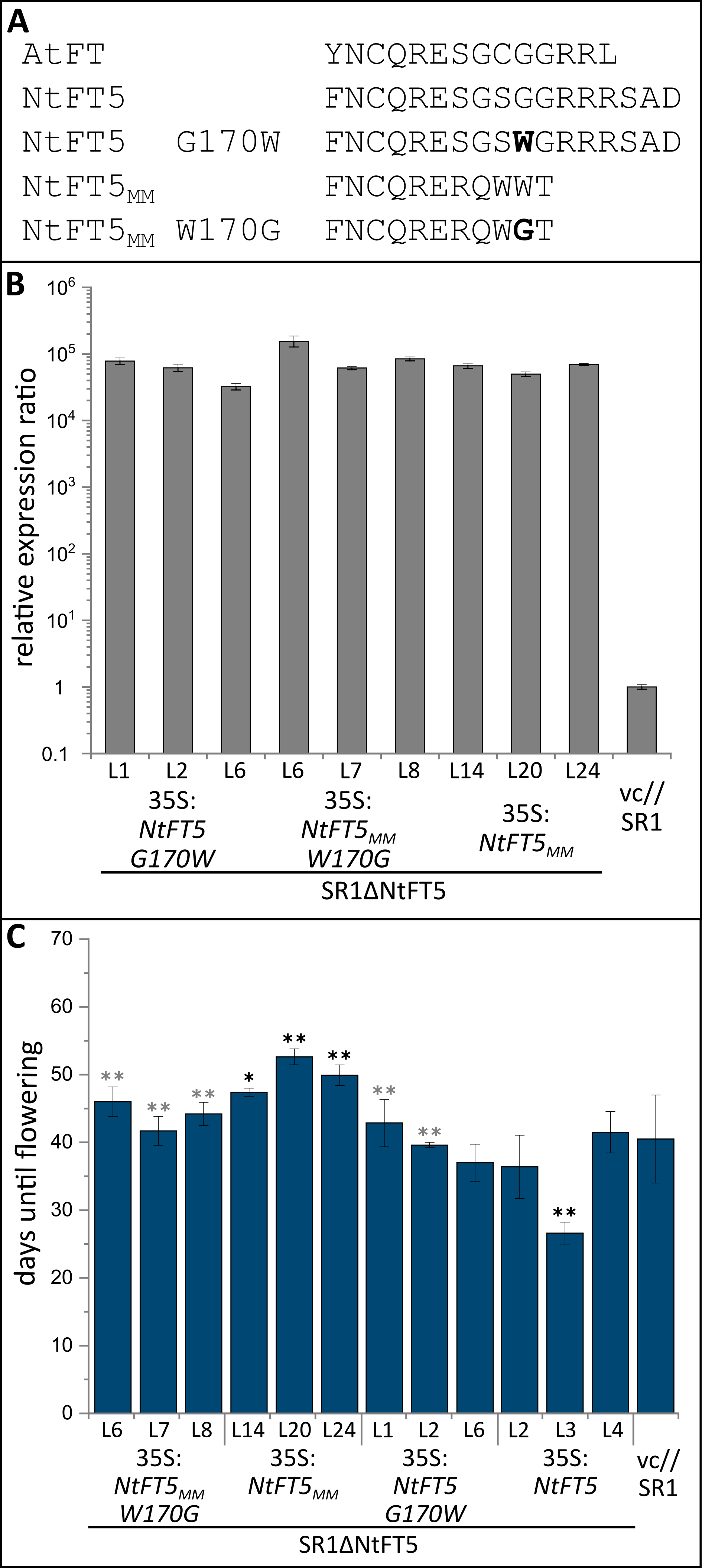

**Figure S6.** **Functional characterization of NtFT5_MM W170G_ and NtFT5_G170W_ mutants.**

(A) Partial amino acid sequence alignment of native NtFT5_(MM)_, mutated NtFT5_(MM)_ and AtFT. Amino acid substitutions are shown in bold.

(B) Expression of native and mutated *NtFT5*_(MM)_ was analyzed by qPCR using samples isolated from five pooled seedlings. The values were normalized to the reference gene *NtEF-1α*. Expression in the vector control was set to 1 and error bars represent the combined standard deviation of the target and reference gene technical replicates. The levels of *NtFT5*_MM_ mRNA were 50,000-fold (L20), 66,000-fold (L14) and 69,000-fold (L24) higher than the vector control. The levels of *NtFT5*_G170W_ mRNA were 32,000-fold (L6), 62,000-fold (L2) and 78,000-fold (L1) higher than the vector control. The levels of *NtFT5*_MM W170G_ mRNA were 61,000-fold (L7), 84,000-fold (L8) and 154,000-fold (L6) higher than the vector control.

(C) The number of days until flowering was defined as the period between seed sowing and the day the first flower opened. Strong overexpression of *NtFT5*_MM_ in SR1∆*NtFT5* plants resulted in delayed flowering in all transgenic lines compared to the SR1 vector control, whereas the overexpression of *NtFT5*_MM W170W_ in SR1∆*NtFT5* plants showed a significantly earlier flowering phenotype than 35:*NtFT5*_MM_//SR1∆*NtFT5* lines. The lines overexpressing *NtFT5* showed an early flowering phenotype (significantly earlier in L3) when compared the SR1 vector control, whereas NtFT5_G170W_ lines flowered significantly later (L1 and L3) than 35:*NtFT5*//SR1∆*NtFT5* lines.

Mean values are shown for three 35S:*NtFT5*_MM_//SR1∆*NtFT5* lines (L14 and L24, n = 10; L20, n = 8), three 35S:*NtFT5*_MM_//SR1∆*NtFT5* lines (L2, L3 and L4, n = 10), three 35S:*NtFT5*_MM W170G_//SR1∆*NtFT5* lines (L6, L7 and L8, n = 10), three 35S:*NtFT5*_G170W_//SR1∆*NtFT5* lines (L1, n = 8; L2 n = 10; L6, n = 6) and one vector control line (n = 10) with error bars representing ± 95% confidence intervals. Normal distribution was confirmed by applying the Kolmogorov-Smirnov test. Statistical significance was determined using a pairwise Welch’s *t*-test with Bonferroni-Holm correction (***P* < 0.01; **P* < 0.05; black = significant *vs* vc; gray = significant *vs* native construct).

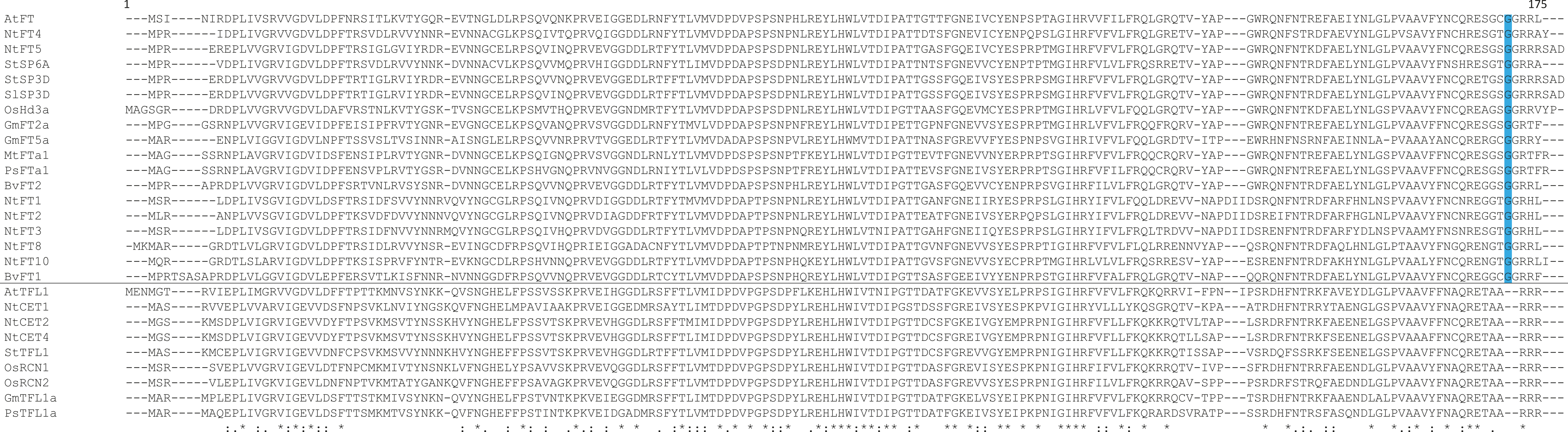

**Figure S7.** **Amino acid sequence alignment of FT-like and TFL-like proteins.**

The glycine residue (highlighted in blue) at position 171 in AtFT is conserved in other FT-like proteins that act as floral activators (green) and repressors (red) in various plant species. TFL-like proteins (black) do not feature a glycine residue at this position. Accession numbers: *Arabidopsis thaliana* AtFT (NM_105222), AtTFL1 (NM_120465) (Kardailsky et al., 1999; Kobayashi et al., 1999); tobacco (*Nicotiana tabacum*) NtFT1 (JX679067), NtFT2 (JX679068), NtFT3 (JX679069), NtFT4 (JX679070), NtFT5 (KY306470), NtFT8 (KY306476), NtFT10 (KY306478), NtCET1 (AF145259), NtCET2 (AF145260), NtCET4 (AF145261) (Amaya *et al*., 1999; Harig *et al.*, 2012; Beinecke *et al.*, 2018); potato (*Solanum tuberosum*) StSP3D (BAV67096), StSP6A (BAV67095), StTFL1 (ABC24691) (Guo et al., 2010; Navarro et al., 2011); tomato (*Solanum lycopersicum*) SlSP3D (AY186735) (Lifschitz et al., 2006); rice (*Oryza sativa*) OsHd3a (NM_001063395), OsRNC1 (AP014967; LOC_Os11g05470.1), OsRNC2 (AP014958; LOC_Os02g32950.1) (Kojima *et al.*, 2002; Nakagawa *et al*., 2002); soybean (*Glycine max*) GmFT2a (AB550122; Glyma16g26660), GmFT5a (AB550126; Glyma_16G044100), GmTFL1a (AB511821) (Kong et al., 2010; Tian et al., 2010); *Medicago truncatula* MtFTa1 (HQ721813) (Laurie et al., 2011); pea (*Pisum sativum*) PsFTa1 (HQ538822) PsTFL1a (AY340579) (Foucher et al., 2003; Hecht et al., 2011); sugar beet (*Beta vulgaris*) BvFT1 (HM448910), BvFT2 (HM448912) (Pin et al., 2010).

**Table S1.** **List of the oligonucleotides used in this study**. Underlined sequences are the overhangs containing the restriction sites used for cloning (lower case letters).

| **Purpose** | **Name** | **5′→3′ sequence** | **Tm** |
| --- | --- | --- | --- |
| pRT104/ pENTR4 cloning | *NtFT5* for Xho I | AGActcgagATGCCAAGAGAACGTGAAC | 55 °C |
|  | *NtFT5* rev Xba I | AGAtctagaTCAATCGGCAGACCTTC |  |
|  | *NtFT5*_FN*_ rev Xba I | ACAtctagaTTAATTAAAATAGACAGCAGCAAC |  |
|  | *NtFT5*_FNC*_ rev Xba I | ACAtctagaTAAACAATTAAAATAGACAGCAGC |  |
|  | *NtFT5*_R61G_ for | GAGATGACCTTgGTACCTTTTACAC |  |
|  | *NtFT5*_R61G_ rev | GTGTAAAAGGTACcAAGGTCATCTC |  |
|  | *NtFT5*_R129A_ for | AGACAATTGGGTgctCAGACAGTG |  |
|  | *NtFT5*_R129A_ rev | CACTGTCTGagcACCCAATTGTCT |  |
|  | NtFT5_MM_ _W170G_ Xba rev | ACAtctagaCTACGTtccCCACTGCCTCTCT |  |
|  | NtFT5_G170W_ Xba rev | ACAtctagaTCAATCGGCAGACCTTCTACGTCCccaACTG |  |
|  | *AtFT* for Sal I | AGAgtcgacATGTCTATAAATATAAGAGACC |  |
|  | *AtFT* rev Not I | AGAgcggccgcCTAAAGTCTTCTTCCTCCGCAG |  |
| P-NtFT5_2.6kb_ | P-NtFT5_2.6kb_ for Xma I | AGAgggcccGTCCGTACCAACTTCTTAGTTC | 57°C |
|  | P-NtFT5_2.6kb_ rev Xho I | AGActcgagGTTGATGCTTAAATAAATAAAACTAAC |  |
| qRT-PCR expression analysis | qRT *NtFT4* for | GATATCCCAGCAACTACAGATACAAG | 67 °C |
|  | qRT *NtFT4* rev | GAAACGGGCAAACCAAGATTGTAAAC |  |
|  | qRT *NtFT5* for | TCCCAAGTTATTAACCAGCCC | 67 °C |
|  | qRT *NtFT5* rev | CATAGCACACAATTTCCTGGC |  |
|  | qRT *NtEF-1a* for | AAGCTGACTGTGCTGTCCTGA | 66.7 °C |
|  | qRT *NtEF-1a* rev | GGTGGTAGCATCCATCTTGTTG |  |
| genomic identification of NtFT7 | NtFT7 for | ACTAGCCATCGTCTTGTGTTTAGCTATTTAG | 56.3 °C |
|  | NtFT7 rev | TGGTAGAGTGATTAAAATGATAAG |  |
| Fragment length analysis | NtFT5 ExIV 396bp for | AATTGGGTCGACAGACAGTG |  |
|  | NtFT5 Stop rev | TCAATCGGCAGACCTTCTAC |  |
| mcBiFC | GGGS Linker NtFT5 for | GGTGGCGGTTCTGGCGGAGGTTCTGGTGGTGGcTCCATGCCAAGAGAACGTG |  |
|  | NtFT5 BsrGI rev | ACAtgtacaTCAATCGGCAGACCTTC |  |
|  | SCYCE_for_Sal | aagagtcgacATGGACAAGCAGAAGAAC |  |
|  | HA_rev_Linker_ BamHI | aagaGGATCCaccaccagaacctccgccagaaccgccaccAGCGTAATCTGGAAC |  |

**Table S2.** **Geno- and phenotype of F_2_ individuals**. M: MM♂xHicks♀; H: Hicks♂xMM♀

| **individual** | **genotype** | **phenotype** | **leaf number** | **days until flowering** |
| --- | --- | --- | --- | --- |
| **M001** | NtFT5/NtFT5_MM_ | flowering | 23 | 56 |
| **M004** | NtFT5/NtFT5_MM_ | flowering | 20 | 56 |
| **M006** | NtFT5/NtFT5_MM_ | flowering | 20 | 60 |
| **M011** | NtFT5/NtFT5_MM_ | flowering | 32 | 77 |
| **M012** | NtFT5/NtFT5_MM_ | flowering | 32 | 58 |
| **M013** | NtFT5/NtFT5_MM_ | flowering | 33 | 67 |
| **M014** | NtFT5/NtFT5_MM_ | flowering | 21 | 56 |
| **M016** | NtFT5/NtFT5_MM_ | flowering | 17 | 61 |
| **M017** | NtFT5/NtFT5_MM_ | flowering | 32 | 67 |
| **M018** | NtFT5/NtFT5_MM_ | flowering | 24 | 56 |
| **M021** | NtFT5/NtFT5_MM_ | flowering | 31 | 61 |
| **M024** | NtFT5/NtFT5_MM_ | flowering | 26 | 48 |
| **M026** | NtFT5/NtFT5_MM_ | flowering | 28 | 50 |
| **M027** | NtFT5/NtFT5_MM_ | flowering | 45 | 95 |
| **M028** | NtFT5/NtFT5_MM_ | flowering | 23 | 57 |
| **M034** | NtFT5/NtFT5_MM_ | flowering | 23 | 50 |
| **M037** | NtFT5/NtFT5_MM_ | flowering | 20 | 49 |
| **M039** | NtFT5/NtFT5_MM_ | flowering | 24 | 55 |
| **M040** | NtFT5/NtFT5_MM_ | flowering | 28 | 57 |
| **M043** | NtFT5/NtFT5_MM_ | flowering | 20 | 46 |
| **M044** | NtFT5/NtFT5_MM_ | flowering | 24 | 56 |
| **M045** | NtFT5/NtFT5_MM_ | flowering | 44 | 105 |
| **M051** | NtFT5/NtFT5_MM_ | flowering | 19 | 62 |
| **M055** | NtFT5/NtFT5_MM_ | flowering | 25 | 53 |
| **M056** | NtFT5/NtFT5_MM_ | flowering | 24 | 55 |
| **M057** | NtFT5/NtFT5_MM_ | flowering | 27 | 53 |
| **M058** | NtFT5/NtFT5_MM_ | flowering | 42 | 105 |
| **M060** | NtFT5/NtFT5_MM_ | flowering | 37 | 73 |
| **M061** | NtFT5/NtFT5_MM_ | flowering | 21 | 48 |
| **M066** | NtFT5/NtFT5_MM_ | flowering | 23 | 53 |
| **M067** | NtFT5/NtFT5_MM_ | flowering | 25 | 56 |
| **M068** | NtFT5/NtFT5_MM_ | flowering | 21 | 53 |
| **M070** | NtFT5/NtFT5_MM_ | flowering | 24 | 53 |
| **M073** | NtFT5/NtFT5_MM_ | flowering | 23 | 49 |
| **M074** | NtFT5/NtFT5_MM_ | flowering | 25 | 56 |
| **M075** | NtFT5/NtFT5_MM_ | flowering | 25 | 57 |
| **M076** | NtFT5/NtFT5_MM_ | flowering | 24 | 73 |
| **M079** | NtFT5/NtFT5_MM_ | flowering | 26 | 56 |
| **M080** | NtFT5/NtFT5_MM_ | flowering | 28 | 78 |
| **M082** | NtFT5/NtFT5_MM_ | flowering | 21 | 50 |
| **M083** | NtFT5/NtFT5_MM_ | flowering | 26 | 53 |
| **M086** | NtFT5/NtFT5_MM_ | flowering | 20 | 53 |
| **M088** | NtFT5/NtFT5_MM_ | flowering | 27 | 53 |
| **M090** | NtFT5/NtFT5_MM_ | flowering | 24 | 56 |
| **M094** | NtFT5/NtFT5_MM_ | flowering | 44 | 86 |
| **M095** | NtFT5/NtFT5_MM_ | flowering | 25 | 56 |
| **M097** | NtFT5/NtFT5_MM_ | flowering | 26 | 53 |
| **M098** | NtFT5/NtFT5_MM_ | flowering | 40 | 86 |
| **M099** | NtFT5/NtFT5_MM_ | flowering | 23 | 50 |
| **M100** | NtFT5/NtFT5_MM_ | flowering | 25 | 55 |
| **H002** | NtFT5/NtFT5_MM_ | flowering | 24 | 53 |
| **H005** | NtFT5/NtFT5_MM_ | flowering | 40 | 72 |
| **H006** | NtFT5/NtFT5_MM_ | flowering | 31 | 61 |
| **H008** | NtFT5/NtFT5_MM_ | flowering | 33 | 86 |
| **H009** | NtFT5/NtFT5_MM_ | flowering | 20 | 59 |
| **H010** | NtFT5/NtFT5_MM_ | flowering | 20 | 50 |
| **H013** | NtFT5/NtFT5_MM_ | flowering | 22 | 54 |
| **H014** | NtFT5/NtFT5_MM_ | flowering | 21 | 50 |
| **H016** | NtFT5/NtFT5_MM_ | flowering | 26 | 56 |
| **H018** | NtFT5/NtFT5_MM_ | flowering | 33 | 59 |
| **H022** | NtFT5/NtFT5_MM_ | flowering | 23 | 52 |
| **H023** | NtFT5/NtFT5_MM_ | flowering | 35 | 77 |
| **H026** | NtFT5/NtFT5_MM_ | flowering | 31 | 71 |
| **H027** | NtFT5/NtFT5_MM_ | flowering | 24 | 57 |
| **H028** | NtFT5/NtFT5_MM_ | flowering | 22 | 51 |
| **H030** | NtFT5/NtFT5_MM_ | flowering | 50 | 90 |
| **H031** | NtFT5/NtFT5_MM_ | flowering | 29 | 57 |
| **H032** | NtFT5/NtFT5_MM_ | flowering | 26 | 51 |
| **H034** | NtFT5/NtFT5_MM_ | flowering | 20 | 50 |
| **H035** | NtFT5/NtFT5_MM_ | flowering | 23 | 56 |
| **H037** | NtFT5/NtFT5_MM_ | flowering | 22 | 60 |
| **H038** | NtFT5/NtFT5_MM_ | flowering | 55 | 90 |
| **H039** | NtFT5/NtFT5_MM_ | flowering | 24 | 59 |
| **H040** | NtFT5/NtFT5_MM_ | flowering | 24 | 53 |
| **H042** | NtFT5/NtFT5_MM_ | flowering | 27 | 48 |
| **H043** | NtFT5/NtFT5_MM_ | flowering | 28 | 60 |
| **H044** | NtFT5/NtFT5_MM_ | flowering | 24 | 56 |
| **H045** | NtFT5/NtFT5_MM_ | flowering | 23 | 56 |
| **H046** | NtFT5/NtFT5_MM_ | flowering | 26 | 57 |
| **H048** | NtFT5/NtFT5_MM_ | flowering | 20 | 48 |
| **H049** | NtFT5/NtFT5_MM_ | flowering | 26 | 56 |
| **H058** | NtFT5/NtFT5_MM_ | flowering | 21 | 48 |
| **H059** | NtFT5/NtFT5_MM_ | flowering | 24 | 56 |
| **H060** | NtFT5/NtFT5_MM_ | flowering | 27 | 53 |
| **H061** | NtFT5/NtFT5_MM_ | flowering | 25 | 60 |
| **H063** | NtFT5/NtFT5_MM_ | flowering | 20 | 47 |
| **H064** | NtFT5/NtFT5_MM_ | flowering | 24 | 56 |
| **H065** | NtFT5/NtFT5_MM_ | flowering | 22 | 53 |
| **H066** | NtFT5/NtFT5_MM_ | flowering | 24 | 60 |
| **H067** | NtFT5/NtFT5_MM_ | flowering | 24 | 51 |
| **H068** | NtFT5/NtFT5_MM_ | flowering | 23 | 56 |
| **H069** | NtFT5/NtFT5_MM_ | flowering | 22 | 48 |
| **H072** | NtFT5/NtFT5_MM_ | flowering | 26 | 56 |
| **H073** | NtFT5/NtFT5_MM_ | flowering | 24 | 56 |
| **H075** | NtFT5/NtFT5_MM_ | flowering | 26 | 62 |
| **H076** | NtFT5/NtFT5_MM_ | flowering | 23 | 61 |
| **H077** | NtFT5/NtFT5_MM_ | flowering | 19 | 50 |
| **H079** | NtFT5/NtFT5_MM_ | flowering | 26 | 52 |
| **H080** | NtFT5/NtFT5_MM_ | flowering | 28 | 51 |
| **H081** | NtFT5/NtFT5_MM_ | flowering | 30 | 63 |
| **H082** | NtFT5/NtFT5_MM_ | flowering | 21 | 47 |
| **H083** | NtFT5/NtFT5_MM_ | flowering | 21 | 52 |
| **H084** | NtFT5/NtFT5_MM_ | flowering | 22 | 56 |
| **H085** | NtFT5/NtFT5_MM_ | flowering | 23 | 56 |
| **H086** | NtFT5/NtFT5_MM_ | flowering | 33 | 71 |
| **H088** | NtFT5/NtFT5_MM_ | flowering | 26 | 56 |
| **H089** | NtFT5/NtFT5_MM_ | flowering | 23 | 49 |
| **H090** | NtFT5/NtFT5_MM_ | flowering | 24 | 50 |
| **H091** | NtFT5/NtFT5_MM_ | flowering | 44 | 108 |
| **H094** | NtFT5/NtFT5_MM_ | flowering | 25 | 55 |
| **H096** | NtFT5/NtFT5_MM_ | flowering | 18 | 47 |
| **H097** | NtFT5/NtFT5_MM_ | flowering | 23 | 48 |
| **M002** | NtFT5_MM_ | non-flowering | 70 |  |
| **M009** | NtFT5_MM_ | non-flowering | 63 |  |
| **M015** | NtFT5_MM_ | non-flowering | 63 |  |
| **M019** | NtFT5_MM_ | non-flowering | 65 |  |
| **M020** | NtFT5_MM_ | non-flowering | 49 |  |
| **M031** | NtFT5_MM_ | non-flowering | 43 |  |
| **M035** | NtFT5_MM_ | non-flowering | 37 |  |
| **M036** | NtFT5_MM_ | non-flowering | 49 |  |
| **M041** | NtFT5_MM_ | non-flowering | 37 |  |
| **M047** | NtFT5_MM_ | non-flowering | 45 |  |
| **M048** | NtFT5_MM_ | non-flowering | 64 |  |
| **M053** | NtFT5_MM_ | non-flowering | 36 |  |
| **M077** | NtFT5_MM_ | non-flowering | 47 |  |
| **M078** | NtFT5_MM_ | non-flowering | 44 |  |
| **M081** | NtFT5_MM_ | non-flowering | 52 |  |
| **M084** | NtFT5_MM_ | non-flowering | 54 |  |
| **M085** | NtFT5_MM_ | non-flowering | 46 |  |
| **M087** | NtFT5_MM_ | non-flowering |  |  |
| **M096** | NtFT5_MM_ | non-flowering | 70 |  |
| **H011** | NtFT5_MM_ | non-flowering | 59 |  |
| **H020** | NtFT5_MM_ | non-flowering | 42 |  |
| **H024** | NtFT5_MM_ | non-flowering | 58 |  |
| **H029** | NtFT5_MM_ | non-flowering | 66 |  |
| **H036** | NtFT5_MM_ | non-flowering | 48 |  |
| **H052** | NtFT5_MM_ | non-flowering | 58 |  |
| **H053** | NtFT5_MM_ | non-flowering | 52 |  |
| **H071** | NtFT5_MM_ | non-flowering | 64 |  |
| **H074** | NtFT5_MM_ | non-flowering | 37 |  |
| **H078** | NtFT5_MM_ | non-flowering | 48 |  |
| **H087** | NtFT5_MM_ | non-flowering | 36 |  |
| **H092** | NtFT5_MM_ | non-flowering | 40 |  |
| **H093** | NtFT5_MM_ | non-flowering | 50 |  |
| **H095** | NtFT5_MM_ | non-flowering | 66 |  |
| **H098** | NtFT5_MM_ | non-flowering | 36 |  |
| **H099** | NtFT5_MM_ | non-flowering | 53 |  |
| **H100** | NtFT5_MM_ | non-flowering | 42 |  |
| **M005** | NtFT5 | flowering | 19 | 53 |
| **M007** | NtFT5 | flowering | 22 | 50 |
| **M010** | NtFT5 | flowering | 23 | 50 |
| **M023** | NtFT5 | flowering | 24 | 62 |
| **M025** | NtFT5 | flowering | 23 | 52 |
| **M029** | NtFT5 | flowering | 21 | 50 |
| **M030** | NtFT5 | flowering | 22 | 47 |
| **M033** | NtFT5 | flowering | 24 | 49 |
| **M038** | NtFT5 | flowering | 38 | 86 |
| **M042** | NtFT5 | flowering | 20 | 46 |
| **M046** | NtFT5 | flowering | 19 | 48 |
| **M049** | NtFT5 | flowering | 22 | 49 |
| **M050** | NtFT5 | flowering | 21 | 49 |
| **M059** | NtFT5 | flowering | 23 | 50 |
| **M062** | NtFT5 | flowering | 26 | 42 |
| **M063** | NtFT5 | flowering | 24 | 47 |
| **M064** | NtFT5 | flowering | 21 | 47 |
| **M065** | NtFT5 | flowering | 23 | 53 |
| **M069** | NtFT5 | flowering | 20 | 60 |
| **M071** | NtFT5 | flowering | 21 | 49 |
| **M072** | NtFT5 | flowering | 23 | 49 |
| **M089** | NtFT5 | flowering | 21 | 50 |
| **M091** | NtFT5 | flowering | 19 | 63 |
| **M092** | NtFT5 | flowering | 22 | 48 |
| **M093** | NtFT5 | flowering | 23 | 49 |
| **H001** | NtFT5 | flowering | 23 | 50 |
| **H003** | NtFT5 | flowering | 21 | 48 |
| **H004** | NtFT5 | flowering | 21 | 53 |
| **H007** | NtFT5 | flowering | 20 | 53 |
| **H012** | NtFT5 | flowering | 22 | 50 |
| **H017** | NtFT5 | flowering | 24 | 51 |
| **H019** | NtFT5 | flowering | 23 | 49 |
| **H021** | NtFT5 | flowering | 29 | 48 |
| **H025** | NtFT5 | flowering | 23 | 47 |
| **H033** | NtFT5 | flowering | 21 | 48 |
| **H050** | NtFT5 | flowering | 21 | 51 |
| **H051** | NtFT5 | flowering | 22 | 48 |
| **H054** | NtFT5 | flowering | 24 | 57 |
| **H056** | NtFT5 | flowering | 26 | 63 |
| **H057** | NtFT5 | flowering | 22 | 49 |
| **H062** | NtFT5 | flowering | 23 | 52 |
| **H070** | NtFT5 | flowering | 21 | 48 |
| **Hicks** |  |  |  |  |
| **1** | NtFT5 | flowering | 21 | 46 |
| **2** |  | flowering | 20 | 46 |
| **3** |  | flowering | 20 | 46 |
| **4** |  | flowering | 20 | 46 |
| **5** |  | flowering | 18 | 46 |
| **6** |  | flowering | 21 | 46 |
| **7** |  | flowering | 20 | 46 |
| **8** |  | flowering | 19 | 46 |
| **9** |  | flowering | 20 | 46 |
| **10** |  | flowering | 21 | 46 |
| **MM** |  |  |  |  |
| **1** | NtFT5_MM_ | non-flowering | 53 |  |
| **2** |  | non-flowering | 62 |  |
| **3** |  | non-flowering | 59 |  |
| **4** |  | non-flowering | 50 |  |
| **5** |  | non-flowering | 52 |  |
| **6** |  | non-flowering | 55 |  |
| **7** |  | non-flowering | 52 |  |
| **8** |  | non-flowering | 53 |  |
| **9** |  | non-flowering | 54 |  |

**Methods S1. Cloning procedures for BiFC.**

To generate a suitable negative control for BiFC, different NtFT5 truncations and amino acid substitutions were generated in regions considered important for the interaction between rice FT and FD (Supplementary Fig. S2; Taoka *et al*., 2011). To generate the *NtFT5*_R61G_ and *NtFT5*_R129A_ constructs, we used overlapping primers to introduce the mutation (Supplementary Table S1) and the *NtFT5* sequence cloned in vector pENTR4 as the template. Parental plasmids were digested with DpnI. Of the many chimeric NtFT5 proteins, only the shortest NtFT5 version we tested (NtFT5_FN*_) did not interact with NtFD1 and was therefore selected as the negative control for BiFC analysis.
